# Exploring midgut expression dynamics: longitudinal transcriptomic analysis of adult female *Amblyomma americanum* midgut and comparative insights with other hard tick species

**DOI:** 10.1101/2024.09.20.614175

**Authors:** Stephen Lu, Lucas C. de Sousa Paula, Jose M.C. Ribeiro, Lucas Tirloni

**Affiliations:** Vector Biology Section, Laboratory of Malaria and Vector Research, National Institute of Allergy and Infectious Diseases, Bethesda, MD, USA; Tick-Pathogen Transmission Unit, Laboratory of Bacteriology, National Institute of Allergy and Infectious Diseases, Hamilton, MT, USA

**Keywords:** ticks, blood meal digestion, midgut, RNA-sequencing

## Abstract

**Background:** Female ticks remain attached to their host for multiple days to complete a blood meal. This prolonged feeding period is accompanied by a significant increase in the tick’s size and body weight, paralleled by noteworthy changes to the tick midgut. While the midgut is recognized for its established role in blood storage and processing, its importance extends to playing a crucial role in the acquisition, survival, and proliferation of pathogens. Despite this, our overall understanding of tick midgut biology is limited.

**Results:** We conducted a comprehensive longitudinal transcriptome analysis of the midgut in adult female *A. americanum* ticks across various feeding stages, including unfed, slow-feeding, and rapid-feeding phases. Our analysis revealed 15,599 putative DNA coding sequences (CDS) classified within 26 functional groups. Dimensional and differential expression analysis highlighted the dynamic transcriptional changes in the tick midgut as feeding progresses, particularly during the initial period of feeding and the transition from the slow-feeding to the rapid-feeding phase. Additionally, we performed an orthology analysis comparing our dataset with midgut transcriptomes from other hard ticks, such as *Ixodes scapularis* and *Rhipicephalus microplus*. This comparison allowed us to identify transcripts commonly expressed during different feeding phases across these three species.

**Conclusion:** Our findings provide a detailed temporal resolution of numerous metabolic pathways in *A. americanum*, emphasizing the dynamic transcriptional changes occurring in the tick midgut throughout the feeding process. Furthermore, we identified conserved transcripts across three different tick species that exhibit similar expression patterns. This knowledge has significant implications for future research aimed at deciphering the physiological pathways relevant within the tick midgut. It also offers potential avenues for developing control methods that target multiple tick species.

## Background

*Amblyomma americanum*, commonly known as the lone star tick, is a three-host tick species prevalent in the eastern, southeastern and south-central regions of the United States (1, 2). The increasing presence of *A. americanum* raises concerns for public health, given its role as a vector for various pathogens. Notably, it has been implicated in the transmission of *Erlichia chafeensis* (3), the causative agent of human monocytic ehrlichiosis, as well as *Francisella tularensis*, responsible for tularemia (4). Additionally, *A. americanum* is associated with the transmission of *Borrelia lonestari* (5), linked to the southern tick-associated rash illness. More recently, this tick species has been proposed as the primary vector for Heartland and Bourbon viruses (6), highlighting its growing significance for public health. Furthermore, bites of *A. americanum* ticks have been associated to the development of alpha-gal syndrome (7).

The feeding cycle of an adult female tick unfolds in three distinct stages. The initial preparatory phase involves the tick attaching to its host’s skin and forming a lesion, setting the stage for the acquisition of its blood meal. The subsequent stage is commonly known as the slow-feeding phase, extending over multiple days. During this phase, the tick experiences a gradual increase in body size and mass as it consumes the host’s blood. The final stage, termed the rapid-feeding phase, occurs in the last 12 to 24 hours of feeding and is characterized by the swift ingestion of host blood (8). Throughout the feeding process, the tick midgut displays notable morphological changes (9), underscoring its high plasticity.

Beyond its established role in blood meal storage and processing (8, 10), the tick midgut serves as the primary entry point for pathogens. Existing evidence supports the idea that specific interactions between ticks and pathogens take place in the midgut, ensuring the survival and proliferation of such pathogens (11, 12). Consequently, different research groups have focused on comprehensively exploring the composition of the tick midgut (13–15) to gain a better understanding of the physiological process within. Furthermore, studies have demonstrated that targeting midgut proteins could be an effective strategy for tick control (16, 17).

It is important to acknowledge that, despite the recent efforts to comprehend tick midgut physiology, the majority of studies have predominantly focused on the early stages of feeding (18, 19). Consequently, these investigations offer a limited perspective on the dynamic changes occurring within the tick midgut throughout the entire feeding process. To bridge this gap, we have previously conducted a comprehensive midgut transcriptome analyses encompassing various feeding stages of *Ixodes scapularis* and *Rhipicephalus microplus* adult female ticks, including the unfed, slow-feeding, rapid-feeding, and post-detachment phases (20, 21). Here, we present an in-depth analysis of the transcriptional alterations observed in the midgut of *A. americanum* adult females during feeding. Recognizing the significance of targeting proteins in the early feeding stages, we compared findings from the *A. americanum* midgut transcriptome with those of *I. scapularis* (20) and *R. microplus* (21). Orthology analyses were performed to identify genes that are commonly expressed in different feeding phases among these tick species. Altogether, this dataset not only enhances our understanding of the adaptations of the tick midgut to the blood meal but also contributes to the identification of potential novel targets for the development of anti-tick control methods.

## Methods

### Ethics statement

Animal experiments were conducted in accordance with the guidelines of the National Institutes of Health on protocols approved by the Rocky Mountain Laboratories Animal Care and Use Committee (2020–065). The Rocky Mountain Veterinary Branch is accredited by the International Association for Assessment and Accreditation of Laboratory Animal Care (AAALAC).

### Tick rearing and midgut dissection

*Amblyomma americanum* ticks were purchased from the tick rearing facility at Oklahoma State University. Unfed ticks were maintained at 21°C and 80-90% relative humidity before infestation. Adult ticks used for midgut extraction were restricted to feed onto the outer part of the ear of four naïve female New Zealand White rabbits with glued orthopedic stockinet. A total of 15 adult females and 15 males (30 ticks per ear, 60 ticks per animal) were placed into the tick containment apparatus and allowed to attach. To group ticks by a blood feeding index, partially fed ticks were collected from host during the feeding, selected based on their engorgement size, and sorted by their average weight in biological triplicates: group unfed (UF, 4.7 ± 0.62 mg, 10 ticks per sample), G1 (6.4 ± 0.60 mg, 5 ticks per sample), G2 (16.4 ± 1.82 mg, 5 ticks per sample), G3 (24.7 ± 3.24 mg, 5 ticks per sample), G4 (67.2 ± 7.30 mg, 5 ticks per sample), G5 (373.9 ± 34.48 mg, 3 ticks per sample), and G6 (577.0 ± 88.50 mg, 3 ticks per sample). After removal from the host, ticks were rinsed with 1% bleach, nuclease-free water, and 70% ethanol, followed by a final rinse with nuclease-free water. Ticks were dissected within two hours after removal from the host. Tick midguts (MGs) were dissected in a fresh ice-cold nuclease-free phosphate-buffered saline (PBS), pH 7.4 (Invitrogen). After dissection, MGs were gently washed in fresh nuclease-free PBS, pH 7.4, containing 4 U/mL of RNAse inhibitor (RNaseOUT, Thermo Fisher Scientific) and a protease inhibitor cocktail (Sigma Aldrich). After washing, dissected MGs were immediately stored in RNAlater (Invitrogen) until total RNA extraction.

### Library preparation, sequencing, and data analysis

Total RNA was isolated using the AllPrep DNA/RNA/Protein mini kit (QIAGEN) according to the manufacturer instructions. RNA integrity and quantification were assessed using a 4200 TapeStation system (Agilent Technologies). The Illumina libraries were constructed using the NEBNextUltraTM II (Directional) RNA with polyA selection library prep kit and sequencing was performed in an Illumina Novaseq 6000 DNA sequencer. The quality of raw Illumina reads were checked using the FastQC tool (https://www.bioinformatics.babraham.ac.uk/projects/fastqc/). Low-quality sequences with a Phred quality score (Q) below 20 and the Illumina adaptors were removed using TrimGalore (https://github.com/FelixKrueger/TrimGalore). Subsequently, reads were merged and *de novo* assembled using Trinity (2.9.0) (22), in single-stranded F mode, and ABySS (2.3.1) (23) with k values ranging from 25 to 95, with increments of 10. The final assemblies were merged, and sequences sharing at least 95% identity were consolidate using the CD-HIT tool (24). The DNA coding sequences (CDS) with an open reading frame (ORF) of at least 150 nucleotides were extracted based on BLASTp results from several databases, including a subset of the non-redundant protein database, the transcriptome shotgun assembly (TSA), and Refseq-invertebrate. The CDS were extracted if they covered at least 70% of a matching protein. Additionally, all ORFs starting with a methionine and with a length of at least 40 amino acids were subjected to the SignalP tool (V3.0). Sequences with a putative signal peptide were mapped to the ORFs, and the most 5’ methionine was selected as the starting point of the transcript (25). Relative quantification of each CDS was performed by mapping the trimmed Illumina reads to the final set of CDS using RSEM (26) and CDS with a TPM ≥ 5 in at least one biological condition was selected for downstream analysis. Functional annotation of the selected CDS was carried out using an *in-house* program that scanned a vocabulary of approximately 450 words and their order of appearance in the protein matches obtained from BLASTp/RPS-BLAST against various databases, including Transcriptome Shotgun Assembly (TSA), a subset from the Non-Redundant (NR), Refseq-invertebrate, Refseq-vertebrate, Refseq-protozoa, UNIPROTKB, CDD, SMART, MEROPS, and PFAM. This annotation process included percent identities and coverage information (27). The final annotated CDS are available for download as a hyperlinked Excel file (Supplementary file 1). Transcriptome completeness was evaluated using the Benchmarking Universal Single-Copy Orthologs (BUSCO) utilizing the Arachnida database as reference (28).

### Statistical analysis

The multidimensional plot and the pairwise differential expression analysis were carried out with the edgeR package (29) for R (30). Statistical significance was considered when LogFoldChange (LogFC) higher than 2 or lesser than -2, alongside a false discovery rate (FDR) less than 0.05 were obtained. The heatmap plot was generated with the pheatmap package using the TPM values and the volcano plots were generated with the ggplot2 package for R. Unsupervised clustering of the filtered CDS were performed with the Expander tool using the CLICK method (31). Ortholog detection was determined by the reciprocal smallest distance (RSD) method (32), run with a coverage ≥ 80% and an e-value ≤ 0.1.

## Results and Discussion

### The temporal transcriptome landscape of *A. americanum* midgut as feeding progress

The Illumina-based RNA-sequencing of the 28 libraries from *A. americanum* midgut at sequential feeding stages (Fig. 1A and B) resulted in 1,068,002,018 high-quality reads. Following our *de novo* assembly and CDS extraction pipeline, we obtained a total of 127,756 putative transcripts. When aligning the trimmed library reads to these sequences, we observed similar mapping rates across all biological conditions (49.5% ± 2.0%, Supplementary table 1). For downstream analysis, we extracted the CDS that exhibited a minimum TPM value of 5 in at least one biological condition, resulting in a final set of 15,599 transcripts. To assess the dataset’s quality, we employed the Benchmarking of Universal Single Copy Orthologs (BUSCO), which indicated similar levels of completeness across all samples (59.9% ± 5.3%, Supplementary table 2). This observation is within range of other transcriptome studies of blood-feeding arthropods (20, 21), and alongside the consistent mapping rates of our libraires, these results underscore the reliability of our dataset, highlighting the absence of significant biases.

**Figure 1:**
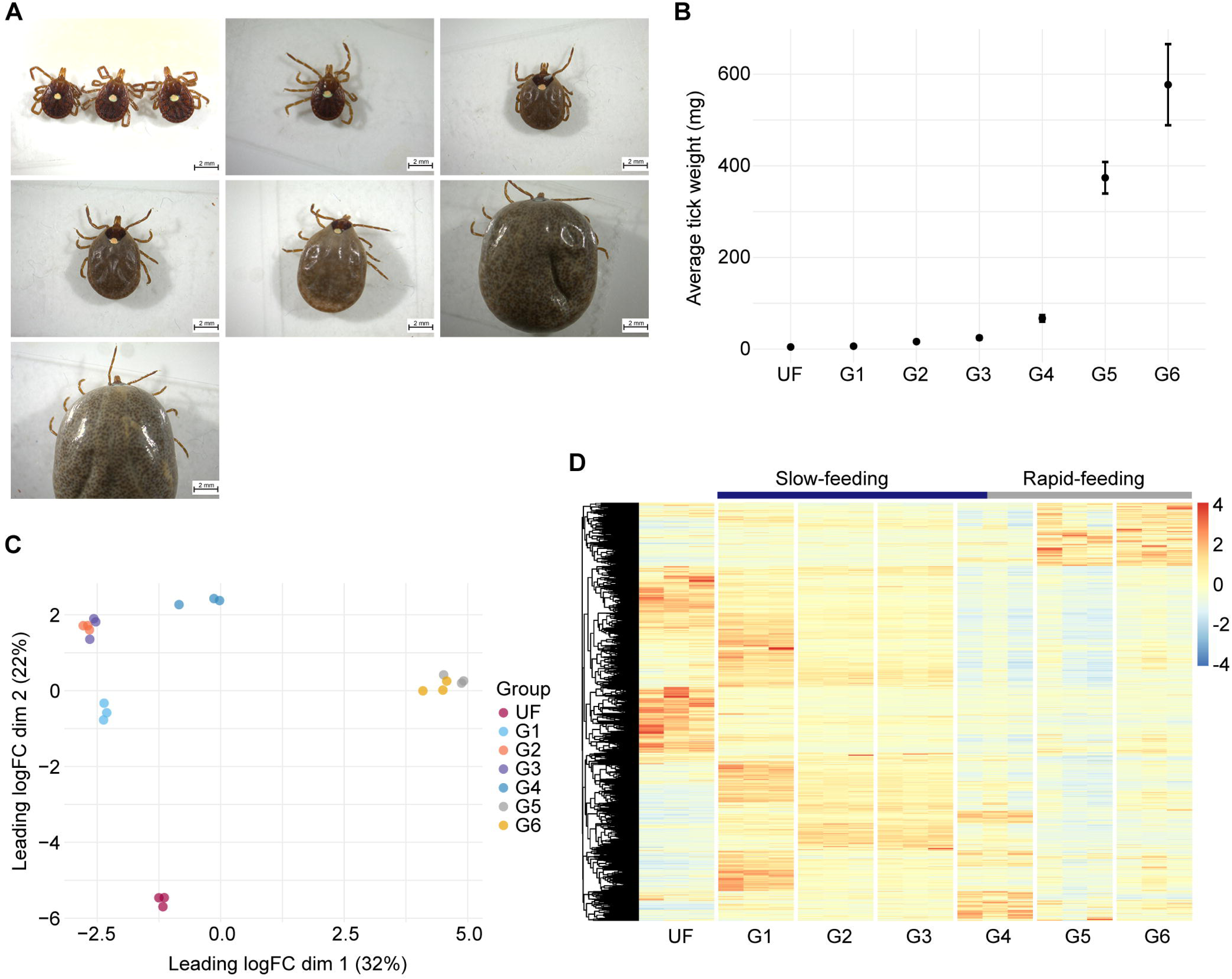
Overview of the transcriptome profile of *Amblyomma americanum* midgut at different feeding stages. **(A)** Representative images of *A. americanum* adult females collected at different feeding stages and their **(B)** average weight (± standard deviation of the mean). **(C)** Multidimension plot of the transcripts identified in *A. americanum* midgut with TPM ≥ 5 in at least one of the biological conditions. **(D)** Heatmap plot of the normalized TPM values of each transcript with TPM ≥ 5 identified in *A. americanum* midgut at each feeding stage.

Dimensional analysis, based on the TPM values from the final 15,599 CDS, revealed that all biological replicates clustered within their respective biological conditions, without any notable outliers (Fig. 1C). Additionally, our samples formed three distinct groups, corresponding to unfed (UF), slow-feeding (G1 – G4), and rapid-feeding (G5 and G6) ticks. A similar pattern was evident when we generated a heatmap plot of the transcripts (Fig. 1D), G4 displaying a transitional transcriptional profile, likely representing the shift from the slow- to the rapid-feeding phases. These results offer valuable insights into the dynamic transcriptional changes occurring in the tick midgut as feeding progresses. Notably, several transcripts that appear to be up-regulated during the initiation of feeding exhibited consistently high abundance throughout the slow-feeding phase. This pattern mirrors recent findings in the midgut of other hard ticks like *I. scapularis* (20) and *R. microplus* (21), suggesting that although the slow-feeding phase spans multiple days, hard ticks in this stage maintain a somewhat conserved transcriptional profile. This observation holds particular interest when considering candidates for developing tick control strategies, since such proteins could remain abundant in the tick midgut during the initial days of feeding, however, further proteomic studies are necessary to validate this hypothesis.

To gain a more comprehensive understanding of the dynamic transcriptional changes occurring in the *A. americanum* midgut during blood acquisition, we performed a pairwise differential expression analyses for each feeding stage compared to the preceding stage (Fig. 2). As reported in other tick species (13, 14, 20, 21), the most significant transcriptional alterations are observed during the transition from unfed ticks (UF) to the initial feeding stage (G1), with 3,477 transcripts showing modulation. In the slow-feeding phase, we noted a moderate number of transcripts displaying differential expression, particularly in the early stages (G2 – G1). Interestingly, despite a 50% increase in weight, there were no discernible differences between the midguts of ticks in the G2 (16.4 mg) and those in the G3 (24.7 mg), indicating nearly identical transcriptional profiles, as suggest by our dimensional analysis (Fig. 1C). Among our comparisons, the G4 – G3 ranked as the third most modulated, involving 1,935 transcripts, while the G5 – G4 comparison was the second most modulated, encompassing 2,374 transcripts. When considering the average weight of ticks in the G3 (24.7 mg), G4 (67.2 mg) and G5 (373.9 mg) groups, alongside the substantial number of differentially expressed transcripts in the G4/G3 and G5/G4 comparisons, it suggests that the G4 group partially represents the end of the slow-feeding phase and the onset of rapid-feeding. Given that the weight difference between G5 and G4 was more than fivefold, future studies aimed at collecting ticks withing the 60 – 380 mg weight range will likely provide a finer resolution of this transition, which represents a key point in the tick life cycle.

**Figure 2:**
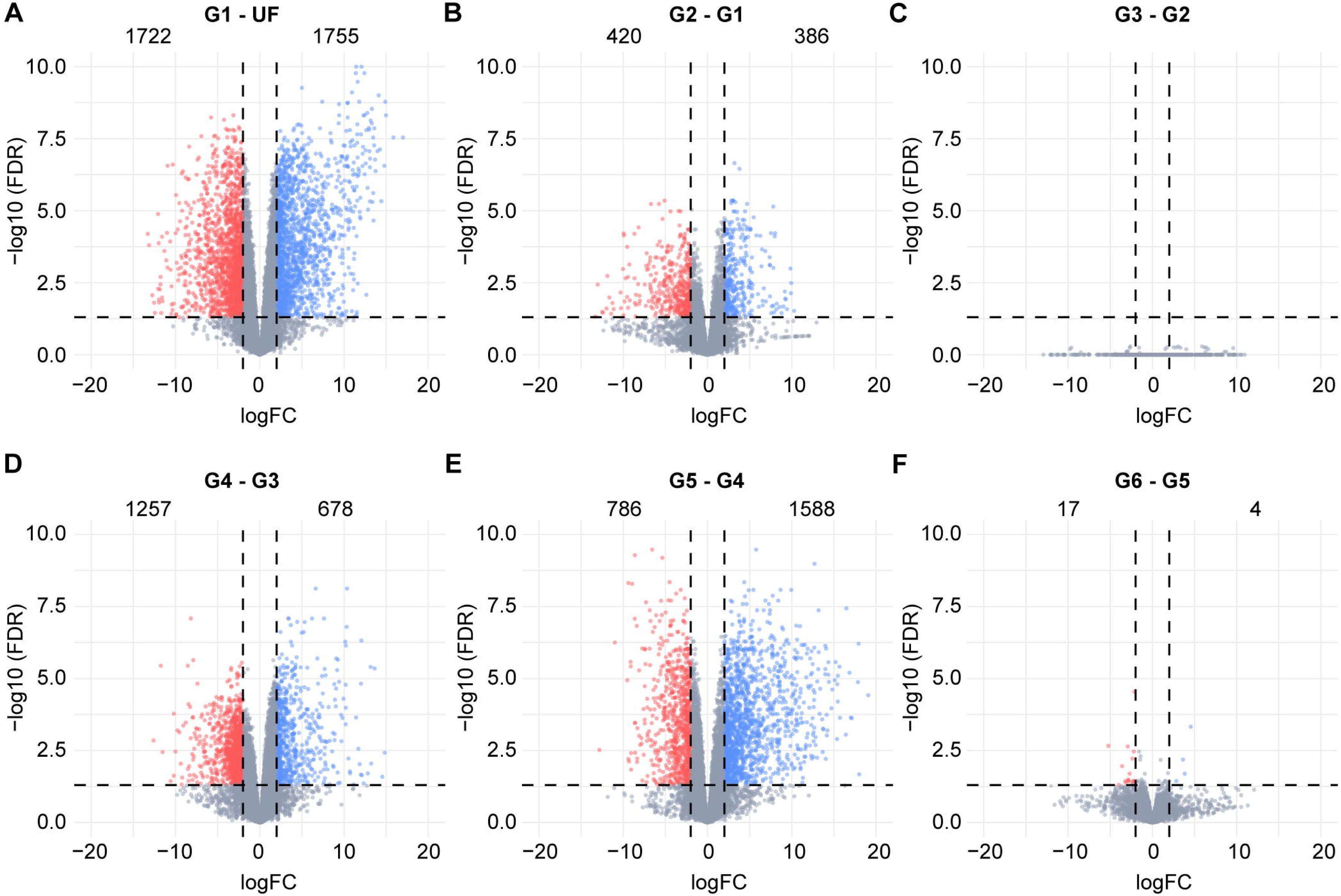
Volcano plots showing the differentially expressed transcripts obtained from the pairwise comparisons between the groups **(A)** G1 and unfed (UF), **(B)** G2 and G1, **(C)** G3 and G2, **(D)** G4 and G3, **(E)** G5 and G4, and **(F)** G6 – G5. The groups G1 – G6 represents ticks in different feeding stages that were group by their average weight; (G1) 6.4 ± 0.60 mg, (G2) 16.4 ± 1.82 mg, (G3) 24.7 ± 3.24 mg, (G4) 67.2 ± 7.30 mg, (G5) 373.9 ± 34.48 mg and (G6) 577.0 ± 88.50 mg. Statistical difference was considered when a transcript presented a LogFC higher than 2 or lesser than -2 (vertical dotted lines), alongside a false discovery rate (FDR) ≤ 0.05 (horizontal dotted lines). Upregulated transcripts are shown as blue dots, downregulated transcripts are shown as red dots and gray dots represents transcripts that were not considered differentially expressed. Numbers inside each plot indicates de number of transcripts differentially expressed.

In parallel to the differential expression analysis, we also conducted an unsupervised clustering of the transcripts based on their TPM values, resulting in the categorization of the putative CDS into six primary patterns (Supplementary figure 1). Cluster 1 comprises 4,466 transcripts predominantly present in the unfed stage. Cluster 2 encompasses, 4,398 CDS, representing sequences highly abundant in the G1 stage, signifying genes promptly induced upon initial contact with host blood, likely playing a crucial role in the early stages of feeding. Congruent with the dimensional analysis and heatmap plots, cluster 3 (2,186 CDS) corresponds to transcripts exhibiting high abundance throughout the tick’s slow-feeding phase. Clusters 4 to 6 consist of CDS mainly abundant during the rapid-feeding phase, but they display distinct patterns, suggesting the presence of specific regulatory mechanism controlling the expression of transcripts during the later feeding stages.

These findings highlight the dynamic transcriptional changes occurring in the midgut of *A. americanum* ticks as they progress through the feeding process. These changes can be categorized into three distinct transcriptional profiles, corresponding to unfed, slow-feeding, and rapid-feeding ticks.

### Functional annotation of *A. americanum* midgut transcripts as feeding progression

To gain a deeper understanding of *A. americanum* midgut physiology during the feeding process, we systematically categorized the 15,999 putative CDS into 26 functional groups. Inspection of these functional classes across different feeding stages provides an overview of the temporal organization of various metabolic processes (Fig. 3). The resulting set of CDS, along with their functional annotation, is available for download in a Windows-compatible hyperlinked Excel file (Supplementary file 1).

**Figure 3:**
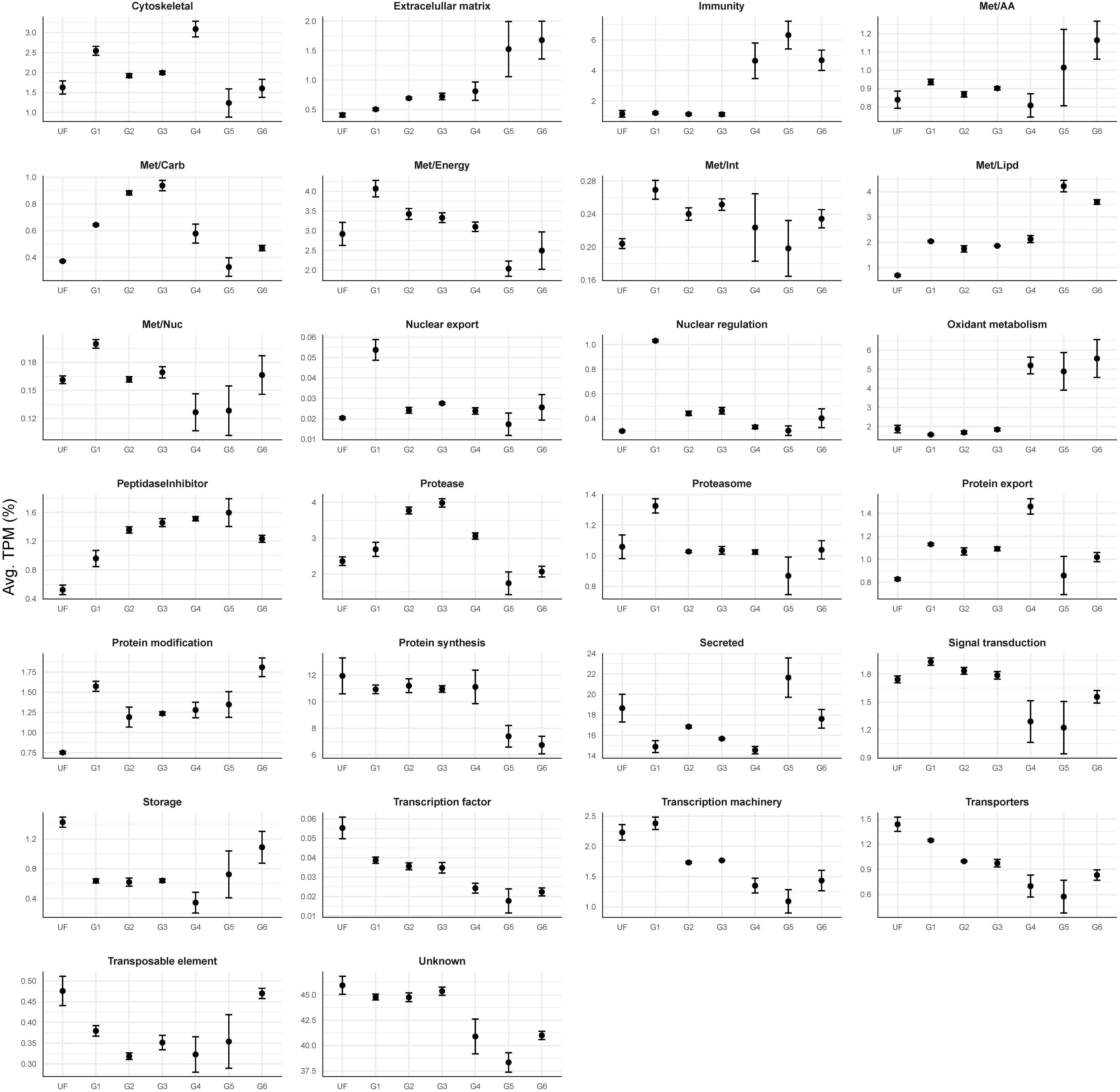
Relative quantification of the 26 functional classes over the different feeding stages of *A. americanum* midgut. Dots represent the average TPM (%) of each class at each biological condition. The error bars represent the standard deviation of the mean.

Notably, our *in-house* classification approach includes the “unknown” class, encompassing transcripts that bear resemblance to deposited sequences of unknown function or exhibit negligible similarities to previously deposited sequences, and thus can be considered potential novel sequences. In the current dataset, this “unknown” functional group was found to be the most abundant in all biological groups, accounting for 38.3% to 45.9% of the total TPM (Fig. 3). The substantial prevalence of this category underscores the existing knowledge gap regarding the composition and function of potential proteins present in the tick midgut, emphasizing the necessity of additional studies focused on this organ. The second most abundant overall functional class was the “secreted”, which encompasses transcripts from different protein families containing a putative secretion signal. Such sequences are commonly reported in the sialome of blood-feeding arthropods and includes sequences bearing similarities to antigen 5-like proteins, lipocalins, lipases, and mucins (33, 34). In the current dataset, the secreted class accounted for 14.6% to 21.6% of all quantified CDS (Fig. 3).

Aside from their abundance, some functional classes presented a very clear pattern as feeding progressed. Specifically, the “immunity” class, which includes multiple transcripts encoding putative antimicrobial peptides such as microplusin-like, lysozyme, and defensins, exhibits low TPM values throughout the unfed (UF) and slow-feeding stages (G1 – G3), representing 1.16% ± 0.03 of the total TPM (Fig. 3). However, a large increase of this class was observed during the rapid-feeding stages (G4: 4.6%, G5: 6.3% and G6: 4.7%). It is interesting to note that this up-regulation of putative antimicrobial transcripts is synchronized with the transition of the slow- to the rapid-feeding phases, in which the tick ingests a substantial volume of blood within a relatively short timeframe (12 – 24 hours) (8). It is likely that the rapid accumulation of the blood meal in the midgut lumen creates a favorable environment for the proliferation of bacteria and other pathogens that could be detrimental to the tick if not properly controlled.

A nearly identical pattern was observed for the “oxidant metabolism” functional group, which encompasses putative catalases, superoxide dismutases, glutathione S-transferases, sulfotransferases, and thioredoxin-like proteins (Supplementary file 1). As the host blood reaches the midgut lumen, red blood cells are lysed through a yet-to-be-discovered mechanism releasing hemoglobin (35). The degradation of hemoglobin leads to the release of significant amounts of heme, which is prone to causing oxidative damage (36). Hence, the rapid increase in the “oxidant metabolism” class during the rapid-feeding phase appears to be a controlled response to the accumulation of heme in the tick midgut.

Interestingly, when comparing the functional classes patterns found in *A. americanum* adult females with those observed in the midgut of *I. ricinus* nymphs (37), we found similar trends. This observation suggests that some of the events occurring in the midgut of nymphs during feeding are conserved during the feeding of adults.

Considering that targeting midgut proteins has been proven to be an effective method of tick control (17, 38), the identification of conserved targets across different tick species and life stages could be of particular interest for developing alternative tick control methods. Hence, in the following sections, we will explore the various transcriptional profiles found in *A. americanum* midgut during blood-acquisition and offer a broad comparison with the midgut of other tick species.

### The midgut of unfed *A. americanum* adult females

To characterize the transcriptional profile of the midgut of unfed ticks, we opted to focus on the 4,466 transcripts that were notably abundant in this stage by our unsupervised clustering analysis (Supplementary figure 1, cluster 1). Based on their TPM values, we found that the predominant functional classes within this cluster were “unknown” (49.1%), “secreted” (20.7%), and “protein synthesis” (8.0%) classes (Fig. 4A).

**Figure 4:**
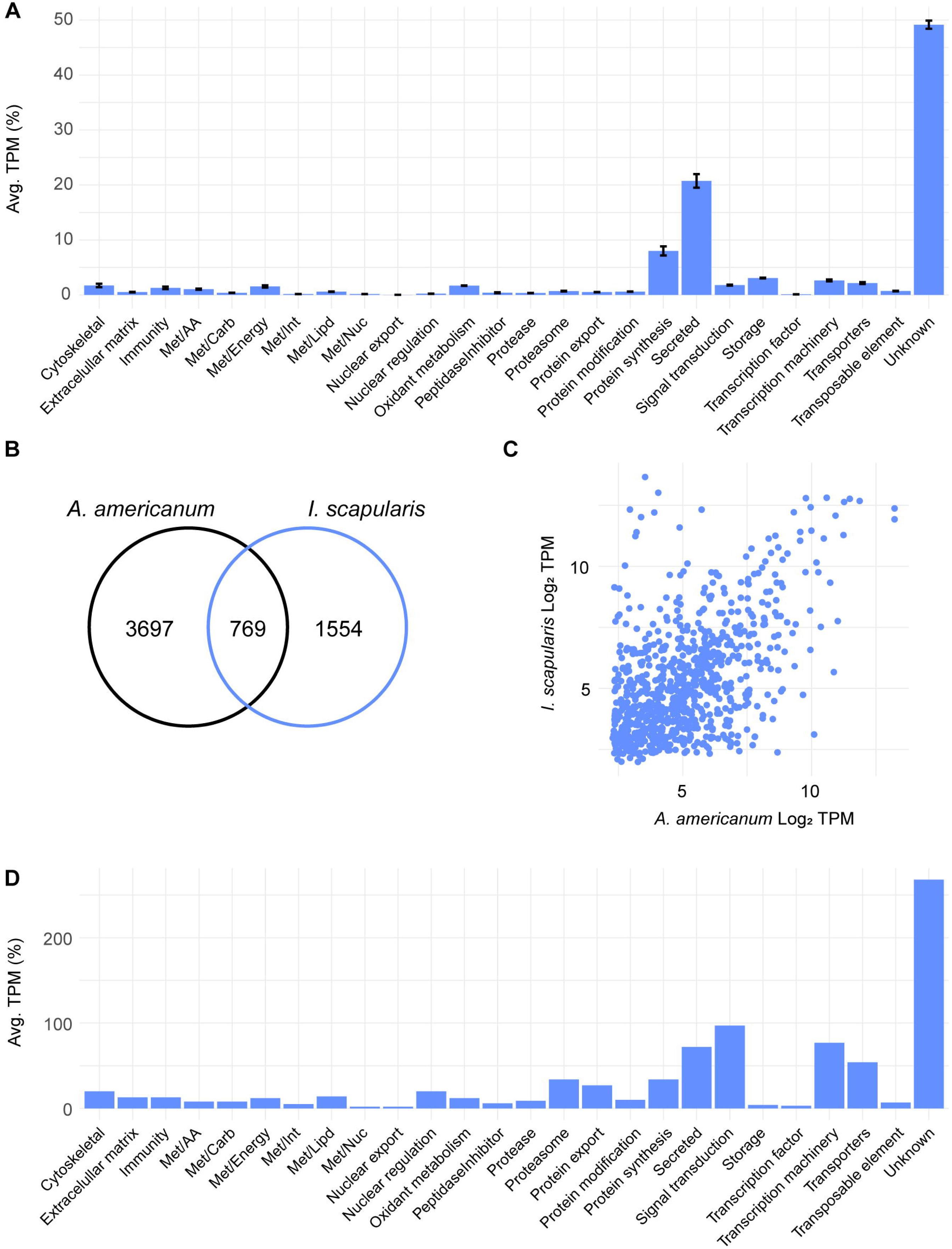
The transcriptional profile of unfed *A. americanum* adult female midgut. (A) Functional classification of 4,466 CDS abundant in the midgut of unfed ticks. Bars represent the average transcript per million (TPM) of each class, the error bars represent the standard deviation of the mean and the number inside each bar indicates the total number of transcripts classified within each class. Comparison between the abundant transcripts found in the midgut of unfed *A. americanum* and *I. scapularis*. (B) Venn diagram represent the number of transcripts unique and orthologous between both ticks. (C) Scatter plot of the Log2TPM of the 769 orthologous transcripts in the midgut of unfed adult females. (D) Functional classification of the shared transcripts between the unfed midguts of *A. americanum* and *I. scapularis* adult females. Bars represent the number of coding sequences (CDS) identified within each class.

Most of the CDS within the “secreted” class that were enriched in this stage were further classified as “unknown”, since they exhibited low or no similarities with previously characterized proteins (Supplementary file 1), underscoring our overall lack of knowledge regarding the potential proteins present in the midgut of *A. americanum* adult females. Furthermore, the abundance of the “protein synthesis” functional class suggests an “active state” of the tick midgut, in which the organ is likely producing or preparing to produce the proteins necessary to accommodate the incoming blood meal. This “preparatory transcriptional profile” was also observed in the midgut of unfed *I. scapularis* adults (20) and unfed *I. ricinus* nymphs (37).

Notably, among the most abundant transcripts in the midgut of unfed females was a putative ferritin (Amseq_49087), with an average TPM of 9,522. Currently, several ferritins have been identified and characterized across different tick species’ midguts (39–41). These can be further classified into type 1 and type 2 ferritins. Type 1 ferritins are intracellular proteins primarily involved in iron storage, with their transcription regulated by the presence of an iron-responsive element (IRE) in their 5’-UTR (42, 43). Conversely, type 2 ferritins lack the IRE domain but possess a putative signal peptide. Further studies indicate that type 2 ferritins are predominantly produced in the tick midgut, facilitating the transport of iron from the midgut to other tissues (44). Given their pivotal role in tick metabolism, these proteins have been suggested as promising targets for anti-tick control strategies (45). Noteworthy is the Amseq_49087 mRNA, which displays an IRE domain within its 5’-UTR and exhibits substantial sequence homology with other type 1 ferritins from ticks (Supplementary figure 2). Although this transcript presents elevated TPM values within the midgut of unfed adult females, its expression remains relatively stable throughout the feeding phase, with TPM values oscillating between 1,147 and 3,633. This consistency suggests the presence and a role of this ferritin throughout the entirety of the adult tick’s feeding cycle.

When comparing the transcriptome profiles of *A. americanum* adult females with those of *I. scapularis* adult females (20), distinct differences emerge. Our unsupervised clustering analysis revealed that *A. americanum* exhibits a greater number of enriched CDS during the unfed stage compared to *I. scapularis* (Fig. 4B). Upon scrutinizing conserved sequences between these ticks at this stage, only 769 CDS met our criteria by RSD analysis (Fig. 4B and Supplementary file 2), underscoring pronounced divergence between the two species. It is important to note that these 769 CDS displayed comparable TPM levels in both tick species (Fig. 4C), exhibiting a Pearson correlation coefficient of 0.52 (p < 0.05). This correlation suggests that these shared sequences are likely present in similar levels within the unfed midgut of both species, however, further proteomic data is necessary to confirm this hypothesis. When exploring their functional classification, the majority of the shared transcripts were classified within the “unknown” (268 CDS), “signal transduction” (97 CDS), “transcription machinery” (77 CDS), and “secreted” (72 CDS) classes (Fig. 4D). Interestingly, when filtering these transcripts based on their overall abundance in the midgut of both ticks, we identified two type 1 ferritins from *I. scapularis* (XP_040076988.1 and XP_029846889.1) with TPM values of 5,304 and 3,890, respectively, ranking them among the most abundant transcripts in unfed ticks. This observation indicates that some transcripts, and potentially metabolic pathways, exhibit the same temporal pattern in the midgut of different tick species, holding significant implications for identifying potential candidates for anti-tick control strategy development, as these conserved pathways may offer heightened efficacy across multiple tick species.

Overall, akin to the kinetic profile observed in the midgut of feeding adult females of *I. scapularis* (20), the midgut of unfed *A. americanum* displayed the most distinct profile compared to other feeding stages (slow- and rapid-feeding ticks, Fig. 1). Specifically, this stage exhibited 4,466 highly enriched CDS, the majority of which are presently categorized as “unknown”. Furthermore, when comparing the current dataset with the abundant transcripts of unfed *I. scapularis* ticks, we identified 769 orthologs shared between both ticks, that presented similar TPM values.

### The midgut of slow-feeding *A. americanum* adult females

As mentioned before, we observed a conserved transcriptional profile during the slow-feeding stage of *A. americanum* adult females, wherein most of the initial changes were induced by the incoming blood meal (Fig. 2).

When exploring the differentially expressed transcripts between the G1 and unfed groups (Table 1), we observed an overall increase of the “lipid metabolism (Met/Lipd)”, “protein modification” and “peptidase inhibitors” functional classes, while the “transcription factor”, “protein synthesis” and “transporters” were the most downregulated classes.

**Table 1:** Functional classification of the differentially expressed transcripts between the midguts of the G1 and unfed *A. americanum* adult females.

| Class | No. of transcripts |  | UF TPM | G1 TPM | G1/UF TPM |
| --- | --- | --- | --- | --- | --- |
|  | Down | Up |  |  |  |
| Nuclear regulation | 6 | 35 | 470.48 | 4056.29 | 8.62 |
| Met/Lipid | 19 | 40 | 1166.13 | 10028.99 | 8.60 |
| Nuclear export | 0 | 9 | 15.33 | 115.21 | 7.52 |
| Protein modification | 7 | 27 | 1828.74 | 6849.27 | 3.75 |
| Extracellular matrix | 5 | 19 | 238.04 | 818.35 | 3.44 |
| Peptidase inhibitors | 21 | 13 | 1940.14 | 5677.87 | 2.93 |
| Met/Carb | 8 | 18 | 257.68 | 751.13 | 2.92 |
| Met/Nuc | 3 | 8 | 124.21 | 240.69 | 1.94 |
| Cytoskeletal | 14 | 19 | 4605.97 | 8456.49 | 1.84 |
| Peptidase | 9 | 16 | 793.32 | 1386.39 | 1.75 |
| Immunity | 24 | 22 | 6179.63 | 6068.01 | 0.98 |
| Transcription machinery | 19 | 19 | 1608.80 | 1475.96 | 0.92 |
| Protein export | 15 | 11 | 895.99 | 755.00 | 0.84 |
| Met/AA | 6 | 13 | 538.60 | 447.54 | 0.83 |
| Met/Energy | 27 | 32 | 2612.86 | 2089.25 | 0.80 |
| Oxidant metabolism | 48 | 36 | 6304.54 | 4801.59 | 0.76 |
| Storage | 3 | 4 | 2064.73 | 1422.31 | 0.69 |
| Signal transduction | 40 | 38 | 1908.36 | 1246.09 | 0.65 |
| Unknown | 956 | 975 | 107006.75 | 59145.66 | 0.55 |
| Proteasome | 15 | 10 | 1072.43 | 526.26 | 0.49 |
| Secreted | 346 | 325 | 75949.14 | 36603.59 | 0.48 |
| Met/Int | 3 | 6 | 479.61 | 226.21 | 0.47 |
| Transposable element | 56 | 11 | 1108.11 | 463.90 | 0.42 |
| Transporters | 59 | 40 | 4079.35 | 1626.14 | 0.40 |
| Protein synthesis | 12 | 9 | 5875.61 | 2149.11 | 0.37 |
| Transcription factor | 1 | 0 | 10.99 | 3.36 | 0.31 |

\*TPM: Transcript Per Million

Currently, our understanding of how ticks metabolize lipids is limited, with only a few studies dedicated to this topic (46, 47). In our current dataset, we identified 59 differentially expressed transcripts between the G1 and unfed ticks (Table 1) that encode putative proteins related to lipid metabolism. Notably, the downregulated CDS included putative enzymes related to lipid catabolism, such as putative acyl-CoA synthetases, which play a crucial role in activating fatty acids prior to β-oxidation, and acyl-CoA oxidases, which catalyze the initial step of fatty acid β-oxidation. Additionally, several transcripts encoding putative triacylglycerol lipases were observed, involved in the hydrolysis of triacylglycerols to glycerol and fatty acids. Conversely, among the upregulated CDS, we identified transcripts for putative enzymes related to lipid biosynthesis. This included enoyl-CoA reductases, important in the synthesis of fatty acids, as well as fatty acid synthetases and sterol O-acyltransferases. The latter encompasses enzymes that convert saturated fatty acids into monosaturated fatty acids, serving as precursors for the synthesis of various lipids. Notably, this trend was also observed within the differentially expressed transcripts of the G2 – G1 comparison (Table 2), where additional triacylglycerol lipases and a putative carnitine O-palmitoyl transferase I, an enzyme related to the transport of long-chain fatty acids into the mitochondria for β-oxidation, were found downregulated.

**Table 2:** Functional classification of the differentially expressed transcripts between the midguts of the G2 and G1 *A. americanum* adult females.

| Class | No. of transcripts |  | G1 TPM | G2 TPM | G2/G1 TPM |
| --- | --- | --- | --- | --- | --- |
|  | Down | Up |  |  |  |
| Met/Carb | 0 | 5 | 317.85 | 1480.93 | 4.66 |
| Extracellular matrix | 3 | 7 | 138.97 | 504.00 | 3.63 |
| Secreted | 91 | 86 | 8596.11 | 30810.94 | 3.58 |
| Storage | 0 | 2 | 8.89 | 30.84 | 3.47 |
| Protein synthesis | 0 | 1 | 1.95 | 6.43 | 3.30 |
| Unknown | 217 | 202 | 13263.17 | 39129.18 | 2.95 |
| Oxidant metabolism | 16 | 7 | 731.77 | 2025.74 | 2.77 |
| Protease | 5 | 4 | 917.89 | 2106.61 | 2.30 |
| Signal transduction | 12 | 6 | 401.03 | 782.21 | 1.95 |
| Met/AA | 4 | 3 | 121.04 | 179.97 | 1.49 |
| Transcription machinery | 2 | 2 | 48.48 | 62.55 | 1.29 |
| Cytoskeletal | 3 | 1 | 55.34 | 62.86 | 1.14 |
| Transposable element | 4 | 8 | 61.37 | 51.40 | 0.84 |
| Met/Lipid | 13 | 10 | 6322.50 | 3808.34 | 0.60 |
| Immunity | 8 | 4 | 1626.83 | 941.31 | 0.58 |
| Met/Energy | 5 | 6 | 185.36 | 90.33 | 0.49 |
| Peptidase inhibitor | 12 | 4 | 439.63 | 188.44 | 0.43 |
| Protein modification | 2 | 1 | 46.36 | 15.93 | 0.34 |
| Transporters | 14 | 4 | 105.39 | 34.47 | 0.33 |
| Proteasome | 1 | 0 | 13.80 | 2.54 | 0.18 |
| Nuclear regulation | 1 | 0 | 1253.03 | 207.92 | 0.17 |
| Met/Int | 1 | 0 | 24.99 | 3.63 | 0.15 |
| Met/Nuc | 2 | 0 | 34.72 | 3.80 | 0.11 |
| Protein export | 3 | 0 | 75.82 | 6.08 | 0.08 |

\*TPM: Transcript Per Million

Transmission electron microscopy of midguts from partially engorged *A. cajennense* adult females revealed the presence of lipid droplets within the digestive cells (9). Although the biochemical characterization of how dietary lipids are digested, absorbed, and transported are still lacking in ticks, based on the transcriptional modulation of CDS encoding putative transcripts related to lipid metabolism, it appears that as the host blood enters the tick midgut and throughout the slow-feeding phase, there is a transcriptional change that switches the lipid landscape from a catabolic profile to an anabolic one, potentially resulting in the formation of lipid droplets. This observation aligns with tick feeding biology, as it has been suggested that during the slow-feeding phase, ticks extract oligonutrients and/or lipids from the blood, while eliminating excess or unnecessary components through saliva or fecal material (48).

The transcripts belonging to the “peptidase inhibitor” class displayed elevated TPM values in the G1 group, exhibiting a 2.93-fold increase compared to the unfed group (Table 1). Within this class, we identified various serine peptidases inhibitors from the serpin, Kunitz-type, and trypsin inhibitor-like (TIL) subfamilies, as well as cystatins, which are tight-binding cysteine peptidase inhibitors. Notably, two CDSs accounted for 89.7% of the total TPM of this class within the G1 group (Supplementary file 1). The CDS Amseq_222115 encoded a putative type-2 cystatin, showing the highest modulation based on TPM values (UF_TPM_ = 528 and G1_TPM_ = 3,906). Additionally, seqSigP-25462, encoding a putative boophilin-like protein, emerged as the second most abundant peptidase inhibitor transcript within the G1 group (TPM = 1190.69). Although other peptidase inhibitors were found differentially expressed between G2 and G1 groups, their overall TPM values were low (Table 2), underscoring the significance of Amseq_222115 and seqSigP-25462 as the primary modulated peptidase inhibitors during the slow-feeding of *A. americanum* adult females.

Currently, several midgut cystatins from ticks have been characterized and associated with the regulation of hemoglobin degradation (49–51), and potentially interacting with cysteine peptidases from tick-borne pathogens (52, 53). Notably, Amseq_222115 exhibited high similarities with other type-2 cystatins that were found to be abundant during the slow-feeding phase of *I. scapularis* (20) and *R. microplus* (21) (Supplementary figure 4). This observation underscores the conserved and important role of cystatins within the tick midgut, in which they likely play similar roles, acting as a major regulator of the activity of tick endogenous cysteine peptidases during the slow-feeding stage.

Boophilin is a serine peptidase inhibitor predominantly found in the midgut of the cattle tick *R. microplus* (54) that contains two Kunitz-type domains, in which the N-terminal domain can bind and inhibit thrombin in a non-canonical manner (55). Furthermore, thrombin-bound boophilin retains the ability to inhibit other serine peptidases, suggesting independent functionality of both Kunitz-type domains. It is proposed that such serine peptidases inhibitors play a crucial role in preventing blood coagulation within the tick midgut, facilitating the accumulation, digestion, and absorption of the blood meal. Similar to boophilin, seqSigP-25462 also exhibits two Kunitz-type domains; however, it possesses two positively charged residues (Arg and Lys) at the P1 of each domain, instead of the Lys and Ala found in boophilin (Supplementary figure 5). This observation suggests that the two domains of seqSigP- 25462 can potentially act as thrombin inhibitors, hence, it is likely that this Kunitz-type inhibitor plays a major role in blocking blood-clotting within the midgut of *A. americanum* adults during the slow-feeding stage.

By comparing the slow-feeding transcriptional profile of *A. americanum* with those observed in *I. scapularis* (20) and *R. microplus* (21), we identified 2,101 and 1,267 shared orthologs, respectively (Fig. 5A). Of those, 768 were common across the three species. Notably, upon evaluating the overall abundance of these shared transcripts within the midgut of each tick species, we found that most exhibited moderate levels of TPM, with a majority presenting Log_2_TPM values ranging from 2 to 10 (Supplementary figure 3A). Additionally, when comparing the abundance of these shared transcripts across the three tick species, we observed comparable levels (Fig. 5B and Supplementary figure 3B).

**Figure 5:**
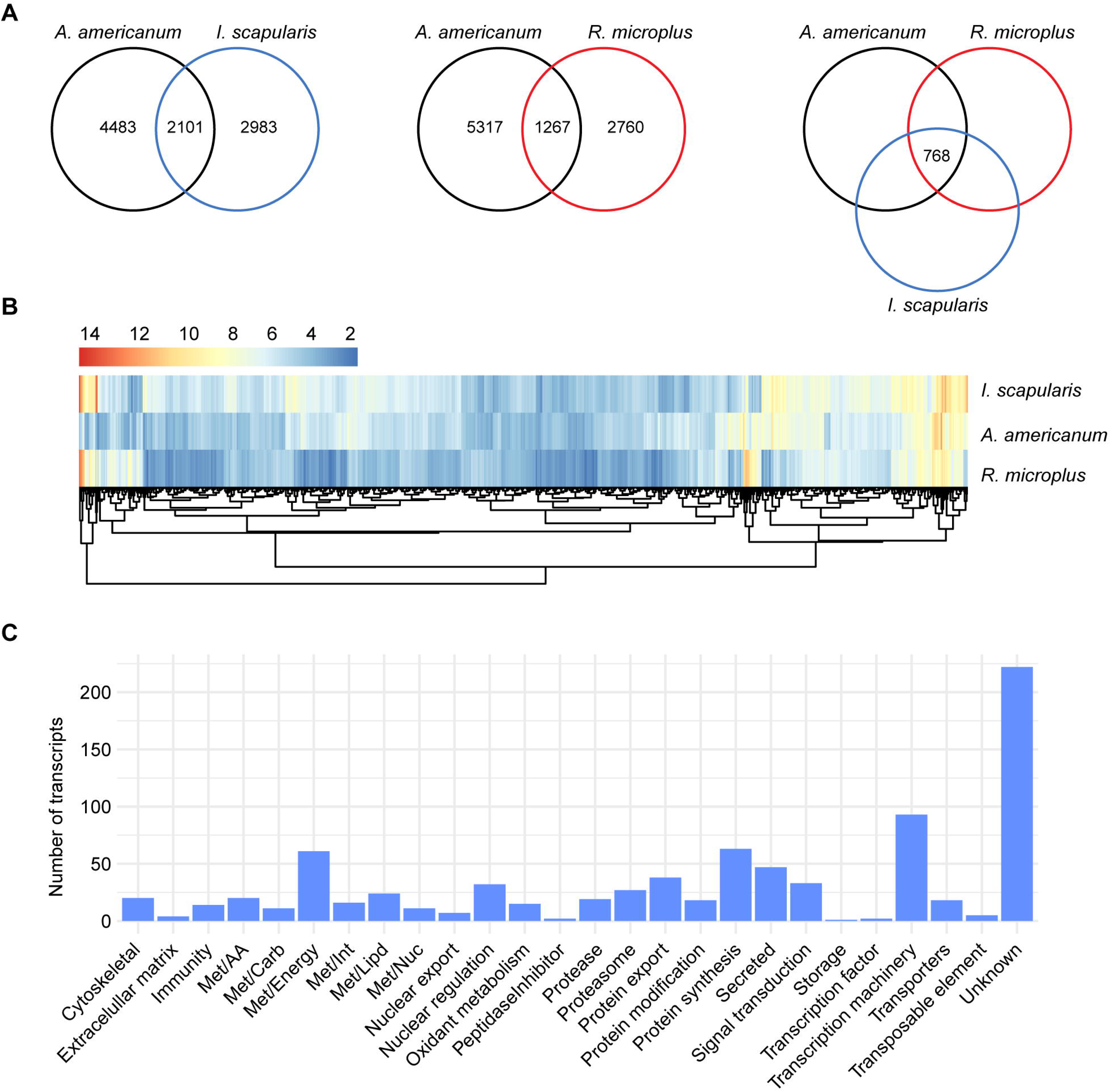
Comparison of the transcriptional profile of the midgut of slow-feeding *A. americanum*, *I. scapularis,* and *R. microplus*. **(A)** Venn diagrams displaying the number of unique and orthologous transcripts across the different tick species. **(B)** Heatmap plot based on the Log_2_TPM of the 768 orthologous transcripts and their **(C)** functional classification.

Further exploration of the functional annotation of the 768 shared transcripts revealed that most were classified as “unknown” (222 CDS). The remaining transcripts fell into functional classes mainly related to housekeeping process, including “transcription machinery” (93 CDS), “protein synthesis” (63 CDS) and “energetic metabolism – Met/Energy” (61 CDS) (Fig. 5C).

Notably, within the “proteases” functional class, we identified conserved transcripts encoding serine peptidases belonging to the S1A subfamily (Supplementary figure 6). While the presence of transcripts coding for putative serine peptidase-like proteins in the tick midgut has been reported previously (13), their exact functions remain unclear. A recent study have demonstrated the absence of trypsin-like proteolytic activity in the midgut of partially fed *I. scapularis* adults, but showing a significant increase post-detachment (56). In *Haemaphysalis longicornis* adult females, various serine peptidase-like proteins were shown to be present within the midgut of partially fed ticks (57, 58). Furthermore, knockdown experiments targeting such transcripts resulted in reduced hemolysis within the tick midgut, suggesting their involvement in blood digestion. It is likely that these peptidases are present as zymogens in the midgut during early feeding days and become activated at later time points. The identification of conserved serine peptidases, particularly abundant during the slow-feeding stage, suggest a similar regulatory mechanism across different tick species, emphasizing the likely importance of these proteins in tick midgut physiology. A role for these proteases in cellular signaling and tissue remodeling, akin to the mechanism observed via protease-activated receptors (PARs) in vertebrates, cannot be ruled out. However, additional studies are needed to ascertain the presence of such receptors activated by proteases in invertebrates. Future studies aimed at the characterization of these serine peptidase-like proteins and their regulation will enhance our current understanding of blood meal digestion in ticks (10).

### The rapid-feeding midgut of *A. americanum* adult females

The final feeding stage of adult females is characterized by the ingestion of a vast amount of host blood within a relatively short timeframe, typically, lasting between 12 to 24 hours. This stage is commonly referred to as the “big sip”, marked by a significant increase in the tick’s body size and mass (Fig. 1A and B). By the conclusion of this stage, the tick naturally detaches from its host. Of particular interest is the observation that the midgut morphology of *Amblyomma* adults during this stage differs noticeably from that of *Ixodes* adults. Specifically, the midgut of *A. cajannense* adult females in this stage exhibits a stratified epithelium with balloon-shaped digestive cells directed towards the midgut lumen (9). In contrast, the midgut of fully engorged *I. ricinus* females features an epidermis forming a cuboidal to squamous epithelium, with stretched flat-like digestive cells (59). This observation illustrates the physiological differences between metastriate and prostriate ticks. Moreover, these differences may also partially explain the unique vectorial capacities of each tick species.

In our dataset, the rapid-feeding stage is predominantly represented by groups 5 and 6 (Fig. 1A and B). However, due to the substantial number of differentially expressed transcripts between G4/G3 and G5/G4, we understand that G4 ticks represent the beginning of the transition from slow- to rapid-feeding ticks. Therefore, we opted to include it in this section. It is important to acknowledge the significant difference in mass between G4 (67.2 ± 7.0 mg) and G5 (373.9 ± 32.5 mg) ticks. Consequently, our differential expression analysis, specifically comparing G5 to G4, partially accounts for the shift from slow- to rapid-feeding ticks. To achieve a more detailed understanding of the changes occurring during this transition, additional efforts focused on collecting partially fed ticks between these groups will be imperative.

In our G4 – G3 differential expression analysis, we uncovered a substantial number of differentially expressed transcripts (Table 3). Notably, the classes exhibiting the most pronounced changes include “immunity” (7.28-fold), “oxidant metabolism” (6.07-fold) and “peptidase inhibitors” (4.63-fold). Within the “immunity” class, we observed an upregulation of numerous transcripts encoding Toll-like receptors, microplusin-like, and lysozyme-like proteins (Supplementary file 1). Remarkably, this elevation of immune-related transcripts seems to persist throughout the rapid-feeding stage of the adult female (Fig. 3 and Table 4). This sustained upregulation likely play an important role in controlling the potential proliferation of microorganism within the midgut lumen, as the blood rapids accumulate. A parallel pattern is evident in the “oxidant metabolism” class (Fig. 3, table 3 and 4), encompassing transcripts likely involved in mitigating oxidative damage caused by the accumulation of heme within the midgut lumen.

**Table 3:** Functional classification of the differentially expressed transcripts between the midguts of the G4 and G3 *A. americanum* adult females.

| Class | No. of transcripts |  | G3 TPM | G4 TPM | G4/G3 TPM |
| --- | --- | --- | --- | --- | --- |
|  | Down | Up |  |  |  |
| Immunity | 10 | 11 | 4944.51 | 35993.81 | 7.28 |
| Oxidant metabolism | 18 | 23 | 5841.90 | 35485.70 | 6.07 |
| Peptidase Inhibitor | 2 | 14 | 1005.71 | 4655.89 | 4.63 |
| Nuclear export | 0 | 2 | 17.07 | 66.44 | 3.89 |
| Protein export | 15 | 14 | 1133.67 | 4007.39 | 3.53 |
| Cytoskeletal | 10 | 14 | 2394.23 | 7356.28 | 3.07 |
| Transposable element | 22 | 13 | 505.5167 | 1340.037 | 2.65 |
| Met/Lipd | 24 | 12 | 4444.32 | 11667.91 | 2.63 |
| Extracellular matrix | 12 | 4 | 632.44 | 1649.09 | 2.61 |
| Secreted | 209 | 136 | 16503.82 | 40711.89 | 2.47 |
| Protease | 7 | 4 | 188.53 | 457.10 | 2.42 |
| Protein modification | 7 | 4 | 459.26 | 1104.46 | 2.40 |
| Met/AA | 7 | 13 | 225.45 | 540.82 | 2.40 |
| Met/Energy | 13 | 7 | 1023.75 | 1929.56 | 1.88 |
| Unknown | 715 | 356 | 24839.76 | 44302.71 | 1.78 |
| Met/Int | 4 | 3 | 150.58 | 213.98 | 1.42 |
| Met/Carb | 9 | 7 | 314.91 | 440.33 | 1.40 |
| Signal transduction | 61 | 12 | 735.47 | 585.38 | 0.80 |
| Transporters | 52 | 15 | 1025.65 | 807.87 | 0.79 |
| Transcription machinery | 36 | 5 | 369.54 | 270.60 | 0.73 |
| Nuclear regulation | 6 | 2 | 34.77 | 17.14 | 0.49 |
| Proteasome | 5 | 3 | 53.58 | 22.56 | 0.42 |
| Met/Nuc | 4 | 1 | 116.74 | 17.06 | 0.15 |
| Protein synthesis | 5 | 0 | 40.94 | 4.97 | 0.12 |
| Transcription factor | 3 | 0 | 37.82 | 4.03 | 0.11 |
| Storage | 3 | 0 | 32.86 | 1.84 | 0.06 |

\*TPM: Transcript Per Million

**Table 4:** Functional classification of the differentially expressed transcripts between the midguts of the G5 and G4 *A. americanum* adult females.

| Class | No. of transcripts |  | G4 TPM | G5 TPM | G5/G4 TPM |
| --- | --- | --- | --- | --- | --- |
|  | Down | Up |  |  |  |
| Met/Lipd | 14 | 33 | 1023.98 | 16770.25 | 16.38 |
| Transcription factor | 0 | 2 | 1.65 | 20.39 | 12.33 |
| Storage | 2 | 3 | 27.09 | 322.79 | 11.92 |
| Extracellular matrix | 18 | 11 | 761.64 | 7883.53 | 10.35 |
| Protein synthesis | 0 | 4 | 2.83 | 16.86 | 5.95 |
| Proteasome | 10 | 2 | 37.17 | 215.04 | 5.79 |
| Met/AA | 12 | 13 | 736.23 | 3964.92 | 5.39 |
| Protein modification | 11 | 10 | 1025.77 | 4808.10 | 4.69 |
| Nuclear regulation | 0 | 2 | 1.88 | 7.69 | 4.10 |
| Immunity | 10 | 24 | 1425.33 | 5198.91 | 3.65 |
| Secreted | 157 | 339 | 35916.35 | 105819.84 | 2.95 |
| Met/Int | 3 | 5 | 129.85 | 350.71 | 2.70 |
| Met/Energy | 8 | 30 | 1202.11 | 2214.43 | 1.84 |
| Unknown | 417 | 831 | 56853.95 | 102607.78 | 1.80 |
| Transcription machinery | 4 | 13 | 250.08 | 349.76 | 1.40 |
| Met/Nuc | 3 | 6 | 77.92 | 106.44 | 1.37 |
| Signal transduction | 17 | 57 | 910.03 | 1204.50 | 1.32 |
| Oxidant metabolism | 27 | 36 | 1613.26 | 1946.96 | 1.21 |
| Transposable element | 8 | 30 | 1330.62 | 1585.53 | 1.19 |
| Transporters | 22 | 49 | 893.36 | 1035.69 | 1.16 |
| Peptidase Inhibitor | 3 | 24 | 10639.80 | 9588.33 | 0.90 |
| Met/Carb | 9 | 8 | 319.46 | 150.12 | 0.47 |
| Protease | 17 | 28 | 17814.66 | 5877.39 | 0.33 |
| Protein export | 9 | 12 | 1826.01 | 455.23 | 0.25 |
| Cytoskeletal | 12 | 8 | 3697.19 | 881.24 | 0.24 |
| Nuclear export | 1 | 0 | 24.28 | 5.45 | 0.22 |

\*TPM: Transcript Per Million

Interestingly, the specific surge of these classes within the midgut of G4 ticks seems to be a controlled response from the tick, preparing itself for the imminent ingestion of vast amounts of host blood during the “big sip”. Furthermore, in *I. scapularis* adult females, a comparable surge in both “immunity” and “oxidant metabolism” classes were also observed. However, this upregulation occurred during the transition from the unfed to slow-feeding ticks and persisted until tick post-detachment (20). In *R. microplus*, the “immunity” class peak occurred during the rapid-feeding stage (21), aligning with *A. americanum*. However, the “oxidant metabolism” class exhibited upregulation during both slow- and rapid-feeding stages. These observations highlight the distinct temporal strategies that each tick species has developed to accommodate their blood meals. Furthermore, they underscore the presence of precise transcriptional regulation mechanisms that are largely unexplored in tick biology.

The comparison between G5 and G4 ticks represents the end of transition from slow- to rapid-feeding adult females. This comparison yielded the second largest number of differentially expressed transcripts (786 downregulated and 1,588 upregulated) within our dataset. Furthermore, when considering the overall TPM values (> 5,000) of the modulated functional classes, we found the “lipid metabolism – Met/Lipd” (16.38-fold), “extracellular matrix” (10.35-fold) and “immunity” (3.65-fold) classes to be the most abundant ones (Table 4).

Notably, within the “Met/Lipd” class, the transcript Amseq_83561 accounted for 56% of the total TPM of this class in G5 ticks (Supplementary file 1). This particular transcript encodes a putative farnesoic acid o-methyltransferases and is presently truncated in its 5’ portion. In insects, this enzyme catalyzes the methylation of farnesoic acid, converting it into methyl farnesoate - a biologically active form of juvenile hormone known for its involvement in various aspects of insect physiology, including molting, reproduction, and vitellogenesis (60). In ticks, studies have suggested that juvenile hormone is not produced (61) and it has no impact on tick vitellogenesis (62, 63). Instead, ticks appear to regulate vitellogenesis through 20-hydroxyecdysone signaling (64, 65). Notably, enzymes belonging to the mevalonate pathway and the juvenile hormone branch have been identified in ticks (66), suggesting their potential in synthesize juvenile hormone precursors. Currently, their specific roles in tick physiology remain unknown.

Similar to the preceding feeding-stages, within the “immunity” class, we identified additional upregulated transcripts encoding putative microplusin-like sequences. Specifically, the CDS Amseq_93759 and seqSigP-254663 accounted for 76.9% of the total TPM of this class (Supplementary figure 7), highlighting the overall abundance of this putative antimicrobial peptide throughout the feeding cycle of *A. americanum* adult females.

Despite the substantial difference in weight between G5 (373.9 ± 32.5 mg) and G6 (577.0 ± 83.4 mg), our differential expression analysis revealed a modest distinction between these two groups (Fig. 3), involving a total of 21 modulated transcripts (17 downregulated and 4 upregulated). This observation underscores a generally consistent transcriptional profile during the rapid-feeding stage of *A. americanum* adult females. In our previous longitudinal transcriptome studies of *I. scapularis* and *R. microplus*, we noted a considerable number of differentially expressed transcripts 24 hours post-detachment (20, 21). However, in the current study, our focus was directed towards exploring the unfed, slow- and rapid-feeding stages of *A. americanum*. We recognize the need for an additional study targeting latter time points to provide a more comprehensive understanding of the transcriptional changes during the feeding cycle of *A. americanum* adult females.

## Conclusions

The feeding process of *A. americanum* adult female ticks is accompanied by a significant morphological transformation characterized by a substantial increase in weight. In this study, we illustrate that the midgut across various feeding stages (unfed, slow-, and rapid-feeding) exhibits distinctive transcriptional profiles, shedding light not only on its composition but also on the presence of precise transcriptional regulatory mechanisms. The dataset presented here serves as a foundational steppingstone for future research endeavors directed at gaining a deeper understanding of tick midgut physiology.

## Authors contributions

Stephen Lu: Conceptualization, Methodology, Formal analysis, Writing – Original draft & editing Lucas Christian de Sousa Paula: Methodology, Reviewer & editing Jose M.C. Ribeiro: Methodology, Formal analysis, Funding acquisition, Writing – Reviewer & editing Lucas Tirloni: Conceptualization, Methodology, Formal analysis, Funding acquisition, Writing – Reviewer & editing

## Acknowledgments

This work utilized the computational resources of the NIH HPC Biowulf cluster (http://hpc.nih.gov). We also thank Rose Perry-Gottschalk, Sj Tudor, and Anita Mora from the Visual and Medical Arts branch (RTB, NIAID, NIH), for figure preparation.

## Data availability

The transcriptome data was deposited to the National Center for Biotechnology Information (NCBI) under BioProject PRJNA1083553 and BioSample accessions SAMN40267879 - SAMN40267899. The raw reads were deposited to the Short Reads Archive of the NCBI under accessions SRR28230933 - SRR28230953. This transcriptome shotgun assembly was deposited at DDBJ/ENA/GenBank under the accession GKSP00000000. The version described in this paper is the first version, GKSP01000000. All supplementary files can be downloaded as a single compressed (.zip) file from the link: https://proj-bip-prod-publicread.s3.amazonaws.com/transcriptome/Aamericanum_Mg_2024/Aamericanum_Mg_SupData.zip

## References

1. Y. P. Springer, L. Eisen, L. Beati, A. M. James, R. J. Eisen, Spatial distribution of counties in the continental United States with records of occurrence of Amblyomma americanum (Ixodida: Ixodidae). J Med Entomol 51, 342–351 (2014).

2. D. E. Sonenshine, Range Expansion of Tick Disease Vectors in North America: Implications for Spread of Tick-Borne Disease. Int J Environ Res Public Health 15 (2018).

3. B. E. Anderson et al., Amblyomma americanum: a potential vector of human ehrlichiosis. Am J Trop Med Hyg 49, 239–244 (1993).

4. C. E. a. D. Hopla, C.M., The Isolation of Bacterium Tularense from the Tick, Amblyomma Americanum. Journal of the Kansas Entomological Society 26, 2 (1953).

5. A. M. James et al., Borrelia lonestari infection after a bite by an Amblyomma americanum tick. J Infect Dis 183, 1810–1814 (2001).

6. A. P. Dupuis, 2nd, R. E. Lange, A. T. Ciota, Emerging tickborne viruses vectored by Amblyomma americanum (Ixodida: Ixodidae): Heartland and Bourbon viruses. J Med Entomol 60, 1183–1196 (2023).

7. S. R. Sharma et al., Alpha-Gal Syndrome: Involvement of Amblyomma americanum alpha-D-Galactosidase and beta-1,4 Galactosyltransferase Enzymes in alpha-Gal Metabolism. Front Cell Infect Microbiol 11, 775371 (2021).

8. D. E. a. R. Sonenshine, R.M., Biology of Ticks (Oxford University Press, ed. 2 ed., 2013), vol. 1.

9. D. Caperucci, G. H. Bechara, M. I. Camargo Mathias, Ultrastructure features of the midgut of the female adult Amblyomma cajennense ticks Fabricius, 1787 (Acari: Ixodidae) in several feeding stages and subjected to three infestations. Micron 41, 710-721 (2010).

10. M. Horn et al., Hemoglobin digestion in blood-feeding ticks: mapping a multipeptidase pathway by functional proteomics. Chem Biol 16, 1053–1063 (2009).

11. U. Pal et al., TROSPA, an Ixodes scapularis receptor for Borrelia burgdorferi. Cell 119, 457–468 (2004).

12. H. Maeda et al., Initial development of Babesia ovata in the tick midgut. Vet Parasitol 233, 39–42 (2017).

13. J. Perner et al., RNA-seq analyses of the midgut from blood- and serum-fed Ixodes ricinus ticks. Sci Rep 6, 36695 (2016).

14. G. A. Landulfo, et al., Gut transcriptome analysis on females of Ornithodoros mimon (Acari: Argasidae) and phylogenetic inference of ticks. Rev Bras Parasitol Vet 26, 185-204 (2017).

15. H. N. S. Moreira et al., A deep insight into the whole transcriptome of midguts, ovaries and salivary glands of the Amblyomma sculptum tick. Parasitol Int 66, 64–73 (2017).

16. K. N. Rand et al., Cloning and expression of a protective antigen from the cattle tick Boophilus microplus. Proc Natl Acad Sci U S A 86, 9657–9661 (1989).

17. P. Willadsen et al., Immunologic control of a parasitic arthropod. Identification of a protective antigen from Boophilus microplus. J Immunol 143, 1346–1351 (1989).

18. C. Stutzer, W. A. van Zyl, N. A. Olivier, S. Richards, C. Maritz-Olivier, Gene expression profiling of adult female tissues in feeding Rhipicephalus microplus cattle ticks. Int J Parasitol 43, 541–554 (2013).

19. A. Oleaga, P. Obolo-Mvoulouga, R. Manzano-Roman, R. Perez-Sanchez, De novo assembly and analysis of midgut transcriptome of the argasid tick Ornithodoros erraticus and identification of genes differentially expressed after blood feeding. Ticks Tick Borne Dis 9, 1537–1554 (2018).

20. S. Lu, L. A. Martins, J. Kotal, J. M. C. Ribeiro, L. Tirloni, A longitudinal transcriptomic analysis from unfed to post-engorgement midguts of adult female Ixodes scapularis. Sci Rep-Uk 13 (2023).

21. S. Lu, J. Waldman, L. F. Parizi, I. Junior, L. Tirloni, A longitudinal transcriptomic analysis of Rhipicephalus microplus midgut upon feeding. Ticks Tick Borne Dis 15, 102304 (2023).

22. M. G. Grabherr et al., Full-length transcriptome assembly from RNA-Seq data without a reference genome. Nat Biotechnol 29, 644–U130 (2011).

23. J. T. Simpson et al., ABySS: a parallel assembler for short read sequence data. Genome Res 19, 1117–1123 (2009).

24. L. Fu, B. Niu, Z. Zhu, S. Wu, W. Li, CD-HIT: accelerated for clustering the next-generation sequencing data. Bioinformatics 28, 3150–3152 (2012).

25. J. D. Bendtsen, H. Nielsen, G. von Heijne, S. Brunak, Improved prediction of signal peptides: SignalP 3.0. J Mol Biol 340, 783–795 (2004).

26. B. Li, C. N. Dewey, RSEM: accurate transcript quantification from RNA-Seq data with or without a reference genome. Bmc Bioinformatics 12 (2011).

27. S. Karim, P. Singh, J. M. C. Ribeiro, A Deep Insight into the Sialotranscriptome of the Gulf Coast Tick, Amblyomma maculatum. Plos One 6 (2011).

28. F. A. Simao, R. M. Waterhouse, P. Ioannidis, E. V. Kriventseva, E. M. Zdobnov, BUSCO: assessing genome assembly and annotation completeness with single-copy orthologs. Bioinformatics 31, 3210–3212 (2015).

29. M. D. Robinson, D. J. McCarthy, G. K. Smyth, edgeR: a Bioconductor package for differential expression analysis of digital gene expression data. Bioinformatics 26, 139–140 (2010).

30. R. C. Team (2021) R: A language and environment for statistical computing. (R Foundation for Statistical Computing, Vienna, Austria).

31. R. Shamir et al., EXPANDER--an integrative program suite for microarray data analysis. BMC Bioinformatics 6, 232 (2005).

32. D. P. Wall, H. B. Fraser, A. E. Hirsh, Detecting putative orthologs. Bioinformatics 19, 1710–1711 (2003).

33. J. M. C. Ribeiro, B. Mans, TickSialoFam (TSFam): A Database That Helps to Classify Tick Salivary Proteins, a Review on Tick Salivary Protein Function and Evolution, With Considerations on the Tick Sialome Switching Phenomenon. Front Cell Infect Mi 10 (2020).

34. L. Tirloni et al., Integrated analysis of sialotranscriptome and sialoproteome of the brown dog tick Rhipicephalus sanguineus (s.l.): Insights into gene expression during blood feeding. J Proteomics 229 (2020).

35. D. Sojka et al., New insights into the machinery of blood digestion by ticks. Trends Parasitol 29, 276–285 (2013).

36. M. Citelli, F. A. Lara, I. D. Vaz, P. L. Oliveira, Oxidative stress impairs heme detoxification in the midgut of the cattle tick, Rhipicephalus (Boophilus) microplus. Mol Biochem Parasit 151, 81–88 (2007).

37. T. Kozelkova et al., Insight Into the Dynamics of the Ixodes ricinus Nymphal Midgut Proteome. Mol Cell Proteomics 22, 100663 (2023).

38. K. N. Rand et al., Cloning and Expression of a Protective Antigen from the Cattle Tick Boophilus- Microplus. P Natl Acad Sci USA 86, 9657–9661 (1989).

39. R. L. Galay et al., Multiple ferritins are vital to successful blood feeding and reproduction of the hard tick. J Exp Biol 216, 1905–1915 (2013).

40. P. Kopácek et al., Molecular cloning, expression and isolation of ferritins from two tick species - Ornithodoros moubata and Ixodes ricinus. Insect Biochem Molec 33, 103–113 (2003).

41. N. W. Githaka et al., Identification and functional analysis of ferritin 2 from the Taiga tick Ixodes persulcatus Schulze. Ticks Tick-Borne Dis 11 (2020).

42. G. Xu, Q. Q. Fang, J. E. Keirans, L. A. Durden, gene coding sequences are conserved among eight hard tick species (Ixodida: Ixodidae). Ann Entomol Soc Am 97, 567–573 (2004).

43. A. M. Thomson, J. T. Rogers, P. J. Leedman, Iron-regulatory proteins, iron-responsive elements and ferritin mRNA translation. Int J Biochem Cell B 31, 1139–1152 (1999).

44. O. Hajdusek et al., Knockdown of proteins involved in iron metabolism limits tick reproduction and development. P Natl Acad Sci USA 106, 1033–1038 (2009).

45. S. Knorr et al., Preliminary Evaluation of Tick Protein Extracts and Recombinant Ferritin 2 as Anti-tick Vaccines Targeting Ixodes ricinus in Cattle. Front Physiol 9 (2018).

46. A. J. Rosendale, M. E. Dunlevy, M. D. McCue, J. B. Benoit, Progressive behavioural, physiological and transcriptomic shifts over the course of prolonged starvation in ticks. Mol Ecol 28, 49–65 (2019).

47. S. Abdullah, S. Davies, R. Wall, Spectrophotometric analysis of lipid used to examine the phenology of the tick Ixodes ricinus. Parasit Vectors 11, 523 (2018).

48. J. M. Ribeiro, The midgut hemolysin of Ixodes dammini (Acari:Ixodidae). J Parasitol 74, 532-537 (1988).

49. K. Yamaji et al., Hlcyst-1 and Hlcyst-2 are potential inhibitors of HlCPL-A in the midgut of the ixodid tick Haemaphysalis longicornis. J Vet Med Sci 72, 599–604 (2010).

50. J. Kotal et al., Mialostatin, a Novel Midgut Cystatin from Ixodes ricinus Ticks: Crystal Structure and Regulation of Host Blood Digestion. Int J Mol Sci 22 (2021).

51. T. H. S. Cardoso, S. Lu, B. R. G. Gonzalez, R. J. S. Torquato, A. S. Tanaka, Characterization of a novel cystatin type 2 from Rhipicephalus microplus midgut. Biochimie 140, 117–121 (2017).

52. S. Lu et al., A novel type 1 cystatin involved in the regulation of Rhipicephalus microplus midgut cysteine proteases. Ticks Tick Borne Dis 11, 101374 (2020).

53. N. Wei et al., A cysteine protease of Babesia microti and its interaction with tick cystatins. Parasitol Res 119, 3013–3022 (2020).

54. T. S. Soares et al., Expression and functional characterization of boophilin, a thrombin inhibitor from Rhipicephalus (Boophilus) microplus midgut. Vet Parasitol 187, 521–528 (2012).

55. S. Macedo-Ribeiro et al., Isolation, cloning and structural characterisation of boophilin, a multifunctional Kunitz-type proteinase inhibitor from the cattle tick. Plos One 3, e1624 (2008).

56. J. Reyes et al., Blood Digestion by Trypsin-Like Serine Proteases in the Replete Lyme Disease Vector Tick, Ixodes scapularis. Insects 11 (2020).

57. T. Miyoshi et al., Molecular and reverse genetic characterization of serine proteinase-induced hemolysis in the midgut of the ixodid tick Haemaphysalis longicornis. J Insect Physiol 53, 195–203 (2007).

58. T. Miyoshi et al., A set of serine proteinase paralogs are required for blood-digestion in the ixodid tick Haemaphysalis longicornis. Parasitol Int 57, 499–505 (2008).

59. J. M. Starck et al., Morphological responses to feeding in ticks (Ixodes ricinus). Zoological Lett 4, 20 (2018).

60. X. Belles, D. Martin, M. D. Piulachs, The mevalonate pathway and the synthesis of juvenile hormone in insects. Annu Rev Entomol 50, 181–199 (2005).

61. P. A. Neese, E. S. D, V. L. Kallapur, C. S. Apperson, R. M. Roe, Absence of insect juvenile hormones in the American dog tick, Dermacentor variabilis (Say) (Acari:Ixodidae), and in Ornithodoros parkeri Cooley (Acari:Argasidae). J Insect Physiol 46, 477-490 (2000).

62. D. Taylor, Y. Chinzei, K. Miura, K. Ando, Vitellogenin Synthesis, Processing and Hormonal- Regulation in the Tick, Ornithodoros-Parkeri (Acari, Argasidae). Insect Biochem 21, 723–733 (1991).

63. Y. Chinzei, D. Taylor, K. Ando, Effects of juvenile hormone and its analogs on vitellogenin synthesis and ovarian development in Ornithodoros moubata (Acari: Argasidae). J Med Entomol 28, 506-513 (1991).

64. A. Seixas, K. J. Friesen, W. R. Kaufman, Effect of 20-hydroxyecdysone and haemolymph on oogenesis in the ixodid tick Amblyomma hebraeum. J Insect Physiol 54, 1175–1183 (2008).

65. K. J. Friesen, W. R. Kaufman, Effects of 20-hydroxyecdysone and other hormones on egg development, and identification of a vitellin-binding protein in the ovary of the tick, Amblyomma hebraeum. J Insect Physiol 50, 519–529 (2004).

66. J. Zhu et al., Mevalonate-Farnesal Biosynthesis in Ticks: Comparative Synganglion Transcriptomics and a New Perspective. Plos One 11, e0141084 (2016).

