## Supplementary data for "Exploring midgut expression dynamics: longitudinal transcriptomic analysis of adult female *Amblyomma americanum* midgut and comparative insights with other hard tick species"

**Supplementary table 1**: Mapping rates of the trimmed Illumina reads to the putative CDS obtained from the *de novo* assembly of *A. americanum* midgut transcripts.

| Sample | Reads not aligned | Reads aligned | Total reads | Reads aligned (%) |
| --- | --- | --- | --- | --- |
| UF.A | 28,710,730 | 26,258,152 | 54,968,882 | 47.76 |
| UF.B | 30,391,122 | 25,576,442 | 55,967,564 | 45.69 |
| UF.C | 27,004,578 | 23,953,610 | 50,958,188 | 47.00 |
| G1.A | 22,386,518 | 20,958,188 | 43,344,706 | 48.35 |
| G1.B | 23,009,394 | 21,885,922 | 44,895,316 | 48.74 |
| G1.C | 24,845,561 | 21,745,039 | 46,590,600 | 46.67 |
| G2.A | 36,856,265 | 33,889,303 | 70,745,568 | 47.90 |
| G2.B | 26,483,883 | 26,363,995 | 52,847,878 | 49.88 |
| G2.C | 27,136,423 | 25,931,353 | 53,067,776 | 48.86 |
| G3.A | 21,878,033 | 20,861,427 | 42,739,460 | 48.81 |
| G3.B | 20,460,881 | 19,995,267 | 40,456,148 | 49.42 |
| G3.C | 21,699,051 | 20,822,443 | 42,521,494 | 48.96 |
| G4.A | 19,853,516 | 22,085,700 | 41,939,216 | 52.66 |
| G4.B | 25,514,147 | 27,272,865 | 52,787,012 | 51.66 |
| G4.C | 22,419,683 | 26,471,151 | 48,890,834 | 54.14 |
| G5.A | 27,697,473 | 28,030,731 | 55,728,204 | 50.29 |
| G5.B | 24,356,043 | 26,071,825 | 50,427,868 | 51.70 |
| G5.C | 29,489,917 | 30,514,159 | 60,004,076 | 50.85 |
| G6.A | 25,198,802 | 25,094,764 | 50,293,566 | 49.89 |
| G6.B | 30,571,143 | 29,736,863 | 60,308,006 | 49.30 |
| G6.C | 23,382,954 | 24,569,078 | 47,952,032 | 51.23 |

**Supplementary table 2:** Benchmarking of Universal Single Copy Orthologs (BUSCO) statistics for *A. americanum* midgut at each feeding stage.

| Sample | Complete (%) | Single (%) | Duplicate (%) | Fragmented (%) | Missing (%) |
| --- | --- | --- | --- | --- | --- |
| UF | 60.6 | 54.9 | 5.7 | 4.9 | 34.5 |
| G1 | 66.9 | 59.8 | 7.1 | 5.2 | 27.9 |
| G2 | 64 | 57.5 | 6.5 | 4.9 | 31.1 |
| G3 | 64.1 | 57.5 | 6.6 | 4.9 | 31 |
| G4 | 52.1 | 46.4 | 5.7 | 4.1 | 43.8 |
| G5 | 52.4 | 47.2 | 5.2 | 4.3 | 43.3 |
| G6 | 59.3 | 53.6 | 5.7 | 4.8 | 35.9 |

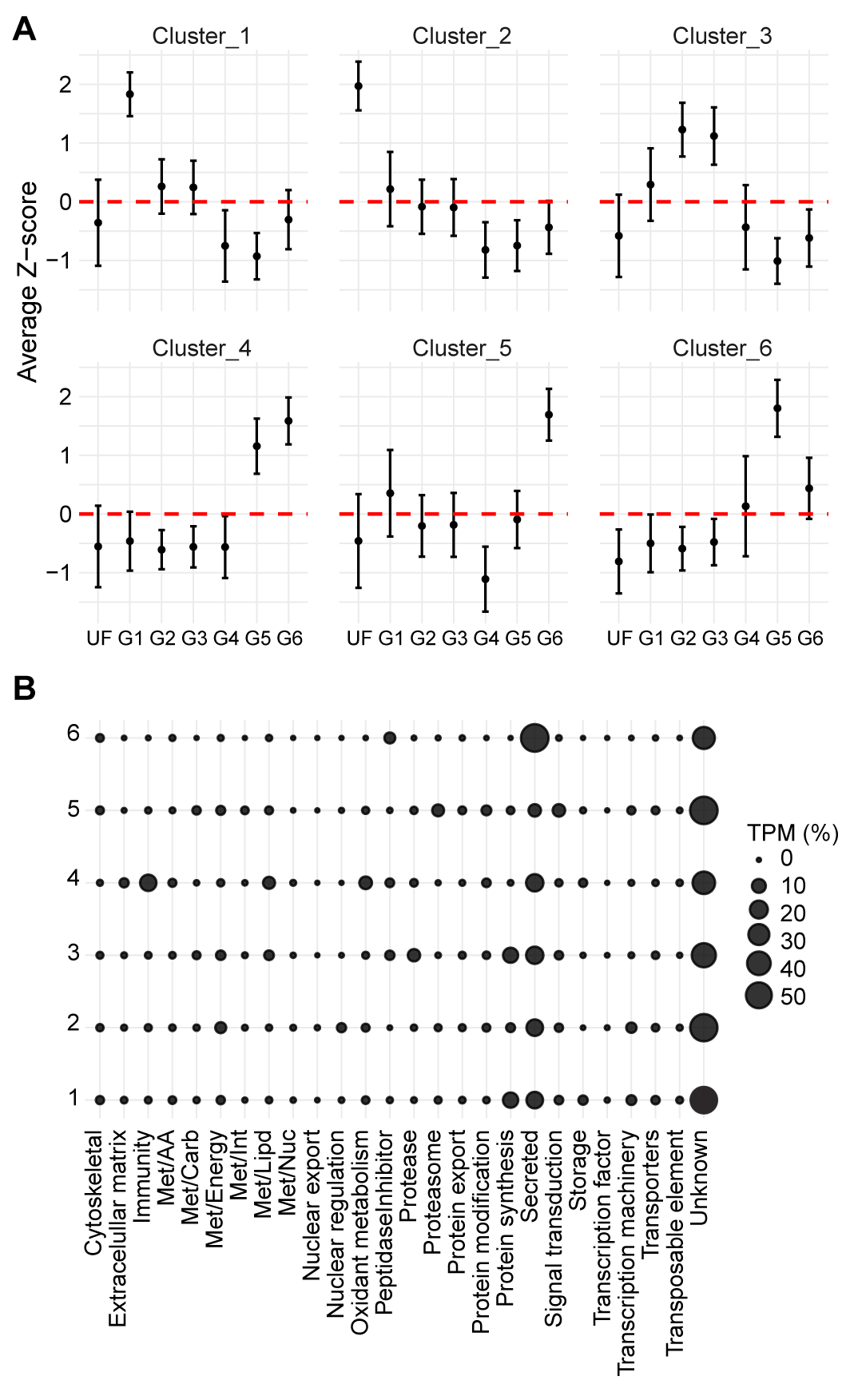

**Supplementary figure 1:** Unsupervised clustering of transcripts identified in *A. americanum* midgut at different feeding stages. **(A)** The dots represent the average Z-score of the transcript per million (TPM) from the transcripts within each cluster and the error bars represent the standard deviation of the mean. The red dotted line marks the zero position in the y-axis. **(B)** Bubble plot representing the abundance of the transcript functional class identified in each cluster.

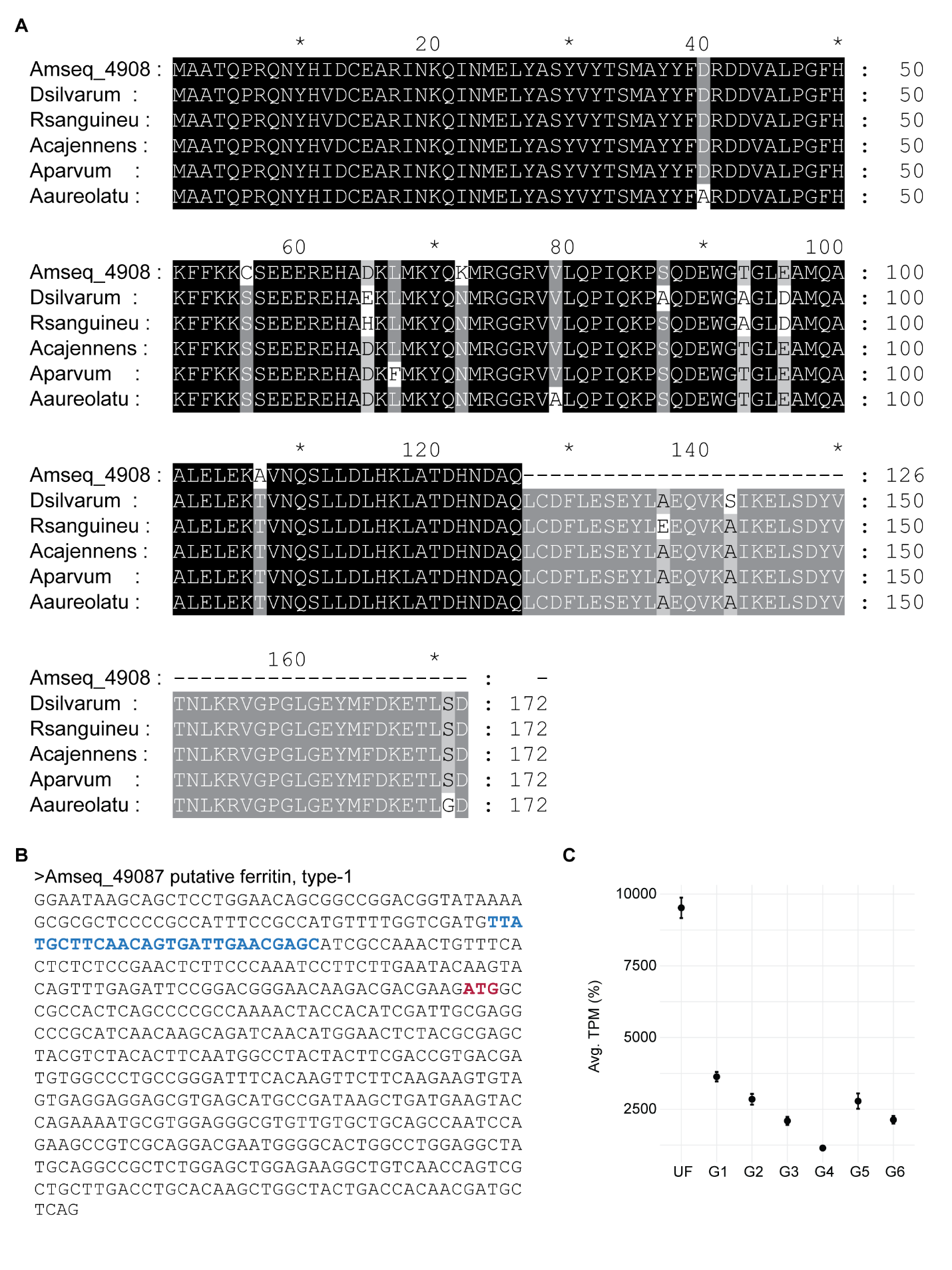
**Supplementary figure 2:** Primary sequence analysis of the putative type 1 ferritin (Amseq_49087) identified in the midgut of unfed adult females *A. americanum* ticks. **(A)** Amino acid alignment of Amseq_49087 with other putative type 1 ferritins from *A. cajennenses* (JAC20482.1), *A. parvum* (JAC25210.1), *A. aureolatum* (JAT93262.1), *R. sanguineus* (AAQ54715.1) and *D. silvarum* (XP_037568710.1). **(B)** Nucleotide sequence of Amseq_49087. The iron-responsive element (IRE) is highlighted in blue and the start codon in red. **(C)** Transcriptional kinetic of Amseq_49087 in the midgut of adult females *A. americanum* ticks at different feeding stages. The dots represent the average transcript per million (TPM) in each biological condition and the error bars represent the standard deviation of the mean.

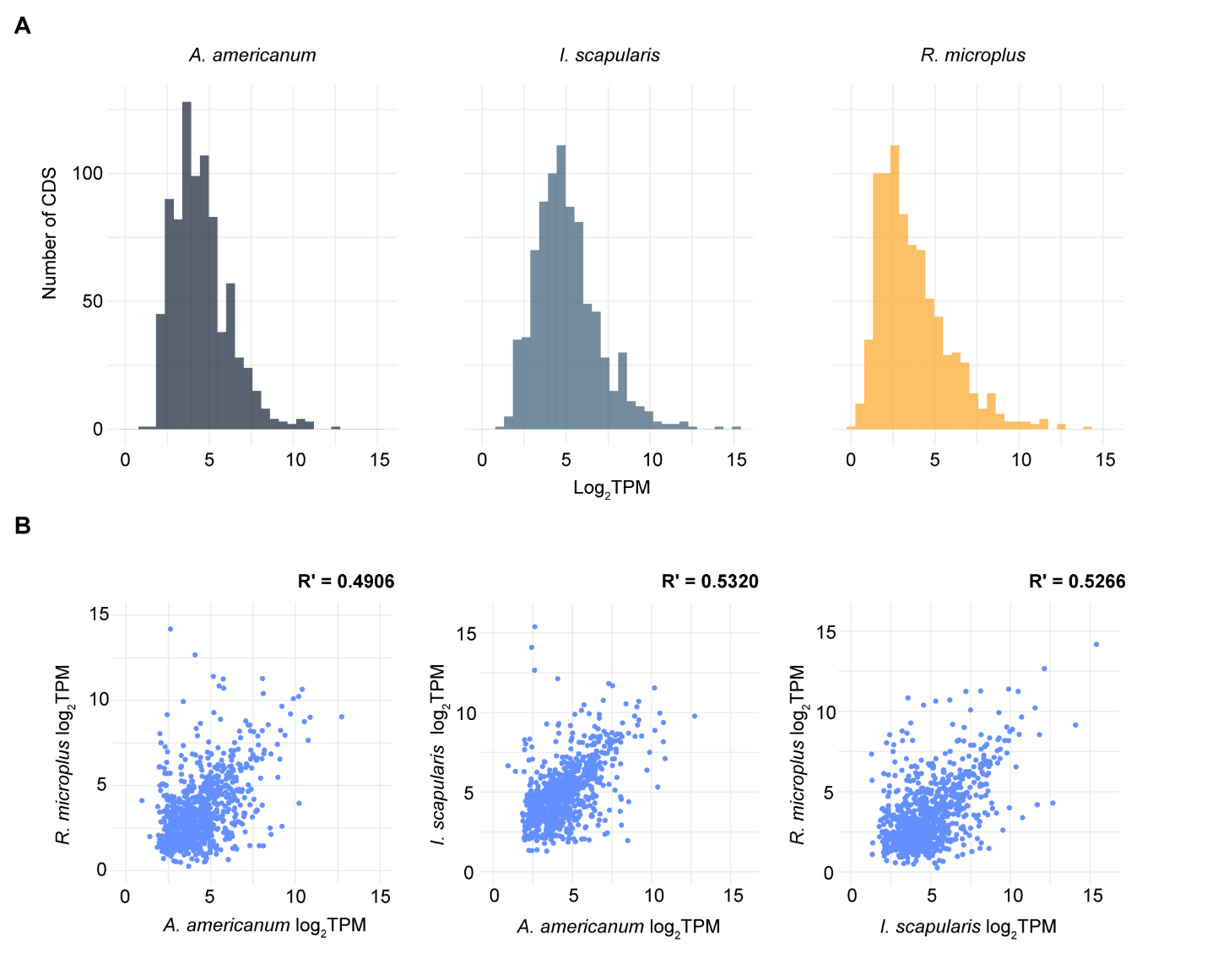
**Supplementary figure 3:** Comparison of the shared transcripts within the midgut of *A. americanum*, *I. scapularis*, and *R. microplus* during the slow-feeding stage. **(A)** Histograms of the 823 shared transcripts among the three tick species based on their Log_2_TPM values. **(B)** Scatter plot of the Log_2_TPM of the 823 shared transcripts in the midgut of slow-feeding ticks. The Pearson correlation coefficient (R^2^) was estimated of each pair-wise comparison.

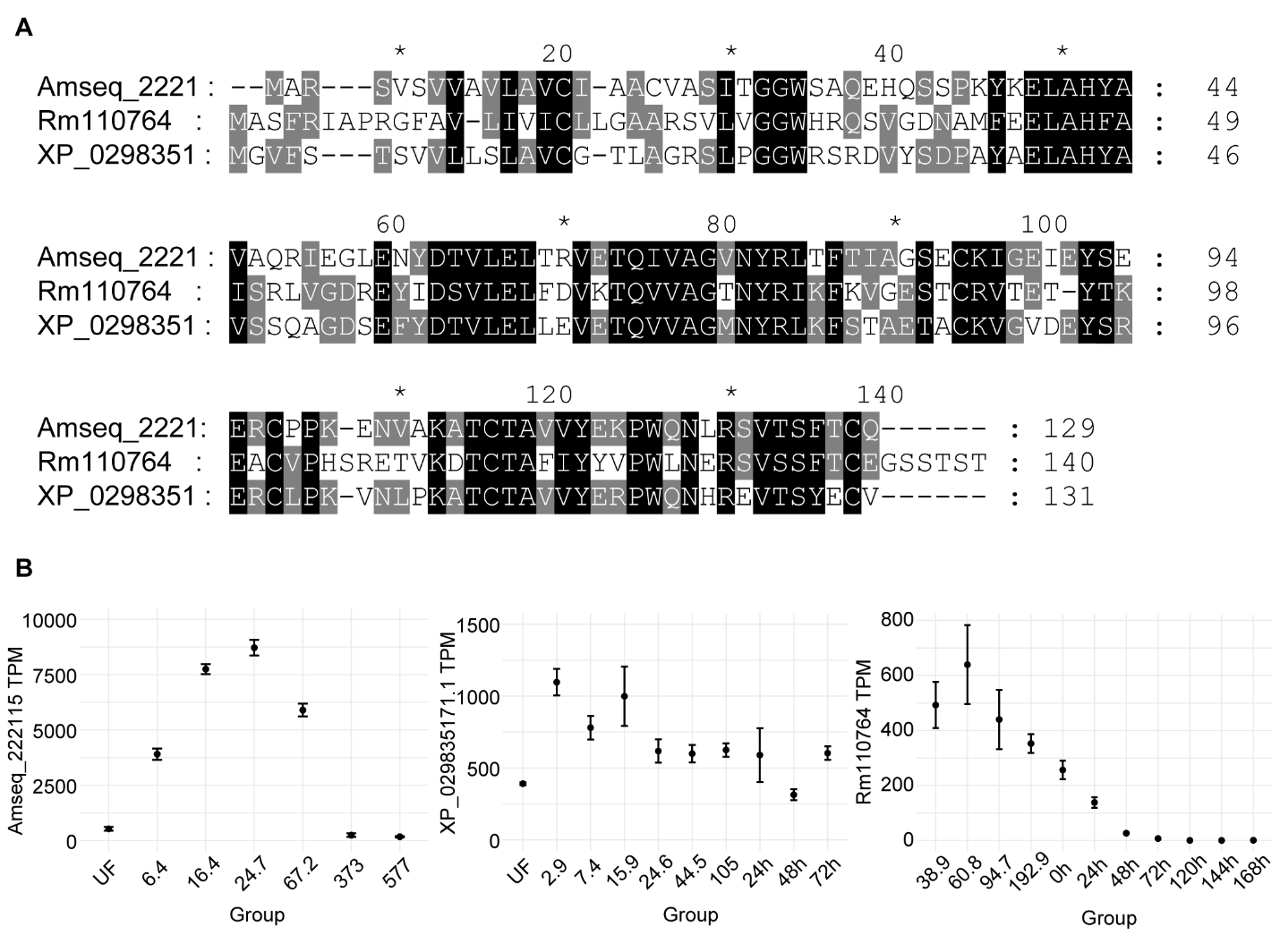

**Supplementary figure 4**: Comparison of putative type-2 cystatins abundant during the slow-feeding phase of *A. americanum*, *I. scapularis* and *R. microplus* adult females. **(A)** amino acid alignment of Amseq_222115 (*A. americanum*), XP_029835171.1 (*I. scapularis*) and Rm110764 (*R. microplus*). Identical (black) and similar (gray) residues are highlighted. (**B)** Transcriptional profile of the three cystatins during the different feeding stages of adult female ticks. Data from XP_029835171.1 and Rm110764 were obtained from [20,21]. Groups represent unfed (UF), partially fed ticks grouped by their average weight (mg) or hours post-detachment (0h – 168h).

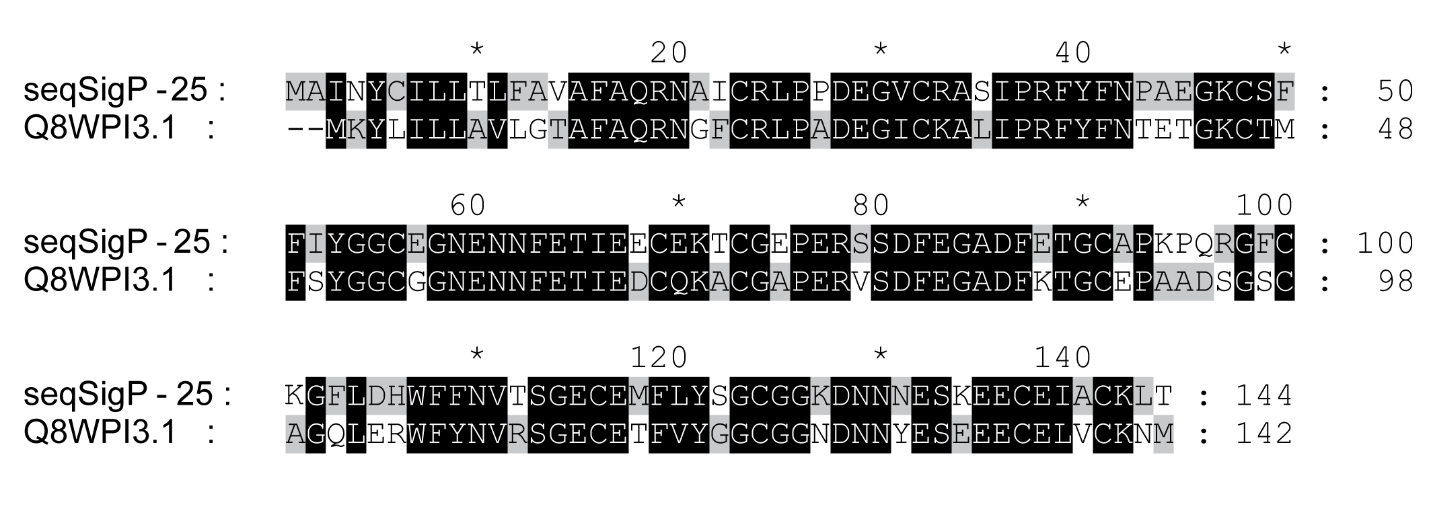
**Supplementary figure 5:** Amino acid alignment of seqSigP-25462 from *A. americanum* and boophilin (Q8WPI3.1) from *R. microplus*. Identical (black) and similar (gray) residues are highlighted.

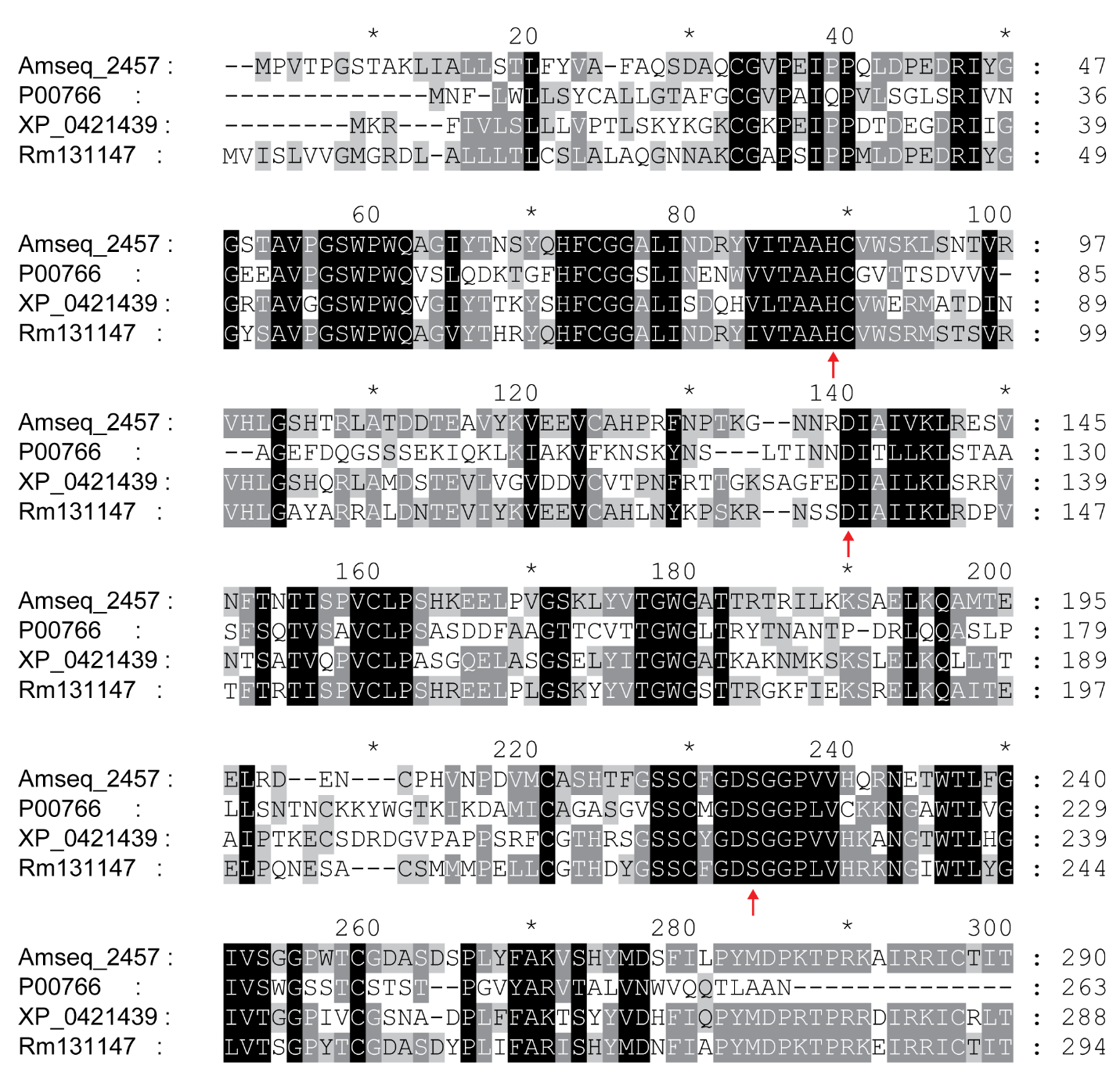
**Supplementary figure 6:** Amino acid alignment of serine peptidases from *Bos taurus* (P00766), *I. scapularis* (XP_042143948.1), *A. americanum* (Amseq_245718) and *R. microplus* (Rm131147). Identical (black) and similar (gray) residues are highlighted, while the putative catalytic residues are indicated by the red arrows.

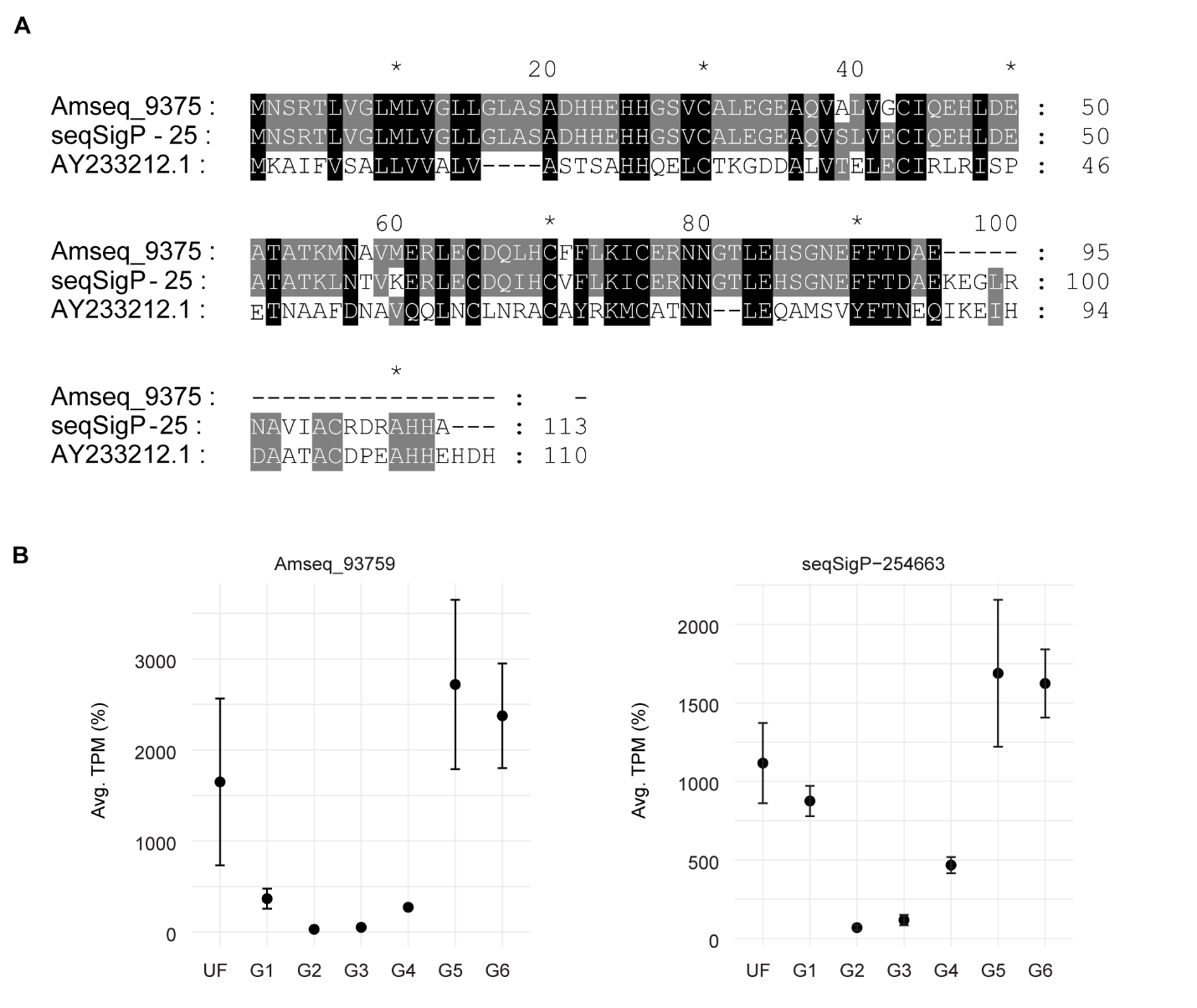

**Supplementary figure 7: (A)** Amino acid alignment of *R. microplus* microplusin (GenBank: AY233212.1) and the predicted amino acid sequence of *A. americanum* microplusin-like transcripts (Amseq_93759 and seqSigP-254663). Identical (black) and similar (gray) residues are highlighted. **(B)** Transcriptional profile of Amseq_93759 and seqSigP-254663. Dots represent the average TPM at each feeding stage, while error bars represent the standard deviation of the mean.

**Supplementary file 1:** Windows-compatible hyperlinked Excel file containing the functional annotation of the 15,999 putative CDS identified in *A. americanum* midgut at different feeding stages.

**Supplementary file 2:** Excel file containing the accession numbers of the shared transcripts within the midgut of unfed and slow-feeding *I. scapularis*, *R. micrplus* and *A. americanum* adult females.
